# Deep FLASH-seq profiling of purified canine sensory neurons uncovers species-specific signatures relevant to pain and itch

**DOI:** 10.64898/2026.04.17.719254

**Authors:** Paula Ledesma Fernandez, Brandi Butler, Heidi Theis, Stefan Paulusch, Elena De Domenico, Greg A Weir, Andrew M Bell

**Affiliations:** Spinal Cord Group, School of Psychology and Neuroscience, University of Glasgow, Glasgow G12 8QQ, United Kingdom; School of Biodiversity, One Health and Veterinary Medicine, University of Glasgow, Glasgow G12 8QQ, United Kingdom; Deutsches Zentrum für Neurodegenerative Erkrankungen (DZNE) e.V., PRECISE Platform for Genomics and Epigenomics at DZNE and University of Bonn, 53127 Bonn, Germany

**Keywords:** Canine, Dorsal root ganglion, single-cell, FLASH-seq, Pain, Itch

## Abstract

Naturally occurring pain and itch disorders in the domestic dog represent an important and underexploited opportunity for translational sensory neuroscience. These conditions largely mirror human disease, highlighting the need for detailed comparative understanding of canine somatosensory neurobiology. Here, we present a single-cell transcriptomic characterisation of the canine dorsal root ganglion (DRG), providing molecular insights into sensory neuron diversity in a species of direct veterinary and biomedical relevance.

We develop a novel mechanical dissociation and fluorescence-activated cell sorting strategy enabling purification of intact whole neurons from adult canine DRG, followed by deep, full-length RNA sequencing using FLASH-seq. This approach yields high-quality transcriptional profiles with molecular depth analogous to deep neuronal profiling in human DRG, enabling resolution of neuronal identities and subtype-specific gene programs.

Using these data, we identify canine sensory neuron clusters conforming to conserved principles of DRG molecular organization observed across species, including peptidergic and non-peptidergic nociceptors, low-threshold mechanoreceptors, proprioceptors, and thermosensory populations. Cross-species comparisons with human and mouse DRG datasets reveal broad conservation of pain- and itch-relevant pathways and therapeutic targets, alongside biologically meaningful divergence.

We further identify species-specific differences in subtype-restricted expression of the pharmacologically relevant receptors IL31RA and SSTR2, which we validate using in situ hybridization and contextualize with human spatial transcriptomic data. Finally, we provide evidence that domestication-associated genes are non-randomly enriched in specific sensory neuron populations, suggesting that evolutionary history may have shaped somatosensory function. These data represent a resource for comparative sensory neuroscience and inform translational interpretation of pain and itch therapeutics across species.

## Introduction

The development of novel therapeutics for pain and itch has been hampered by a marked failure to translate promising pre-clinical analgesic candidates from rodent models[22,66]. Because naturally occurring disease in the pet dog mirrors many aspects of the human condition with greater validity than rodent preclinical models, it has been suggested that studying spontaneous painful disease in companion animals can facilitate the development of therapies for humans while also enhancing veterinary care[2,46]. Pain and itch in dogs are, in many respects, highly similar to pain and itch in humans[37,44,51]. In both species these debilitating sensations arise from naturally occurring conditions such as osteoarthritis and atopic dermatitis. In the domestic dog, these conditions are common, with estimated prevalences of 20% [51] and 3-15% [26], respectively.

Beyond face validity, canine diseases also provide several advantages for translational research. Time courses more closely resemble human chronicity, typically persisting for months or years rather than the shorter durations of many rodent models. Companion animals share important environmental exposures with humans, and many major treatment modalities are common to both species, with comparable pharmacokinetics and clinically relevant dosing intervals. In addition, pain and itch can be measured using a broad suite of outcome tools that extend beyond reflex-based hypersensitivity[36,45], including validated owner-completed behavioural tools, activity and sleep metrics, and quality-of-life instruments[18,50]. However, to use the dog successfully as a translational model requires an in-depth understanding of the fundamental similarities and differences in the molecular basis of somatosensation and pain.

The ability to detect pain- and itch-inducing stimuli in the periphery depends on the existence of multiple classes of primary sensory neurons, each tuned to specific stimuli. Axons of these neurons relay this information to the dorsal horn of the spinal cord via their cell bodies in the dorsal root ganglia (DRG)[7,78]. Classically, the extensive functional heterogeneity of sensory neuron subtypes has been described in terms of cell body size, degree of myelination and electrophysiological properties[17,73]. However, in recent years, single-cell transcriptomics has revolutionised our understanding of the molecular basis of sensory neuron function. Across rodent[34,52,59,62,63,68,79,87], primate[42,49] and human datasets[10,54,58,76,86], single-cell atlases resolve conserved DRG neuron classes and a shared nociceptor taxonomy, alongside species-specific adaptations across subtypes[11,42].

Sensory neurons are compelling therapeutic targets[9], with some novel approaches already used commonly or in development (e.g., monoclonal antibodies targeting NGF[13,27] and IL-31[65,80] signalling, and inhibitors of the Nav1.8 sodium channel) and growing interest in nociceptor-targeted chemogenetic approaches[9,61,84]. However, despite rapid advances in our understanding of rodent and human sensory neuron biology, the cell-level molecular architecture of the canine DRG (cDRG) remains comparatively under-defined[38]. This gap limits both the rational deployment of emerging therapies in veterinary patients and the value of the dog as a translational model for chronic pain and itch.

Here, we use a novel cell-sorting technique combined with FLASH-seq [32] to generate a single-cell transcriptomic atlas of cDRG neurons, and use these data to describe canine sensory neuron populations and quantify cross-species similarity, thereby establishing a framework to inform translation of pain and itch therapeutics.

## Methods

### DRG Collection

All experiments were approved by the Ethics committee of the School of Biodiversity, One Health and Veterinary Medicine, University of Glasgow (ref EA46/24). Tissue was acquired from two groups of dogs. Non-beagle dogs (SSPCA, Glasgow, UK) were euthanized for behavioural reasons; beagle dogs (Charles River Laboratories, Tranent, UK) were part of control groups in research studies unrelated to pain.

Dogs were humanely euthanised with an overdose of barbiturates, and DRGs were collected from the lumbar area within 2 hours from death. DRGs collected for single-cell sequencing and *in situ* hybridization were immediately flash frozen on dry ice and stored at −80°C. DRGs used for immunohistochemistry were immersion fixed in 4% Paraformaldehyde for 48 hours, cryoprotected in 30% sucrose and stored at −150°C.

### Immunohistochemistry

Ganglia were trimmed to remove adjacent nerve roots and connective tissue and embedded in 4% agar. 60 µm- thick transverse sections were cut with a vibrating blade microtome (Leica VT1200, Leica Microsystems, Germany) and non-sequential (1:2) sections were used for downstream experiments. Immunohistochemical reactions were performed on free-floating sections. To measure neuronal diameter, NeuN antibody (clone EPR12763, Catalog N° ab190565; Abcam, UK) was used for neuronal cytoplasm and nuclei identification. Use of this antibody has not been previously reported in canine dorsal root ganglia; however, the same clone is manufacturer-validated for canine brain and spinal cord tissues, is listed as reactive in dog in immunohistochemical applications, and recognises neurons in canine cortex[8]. Sections were incubated overnight with conjugated antibody NeuN-647 (rabbit, 1:1000) at 4°C in Phosphate Buffered saline (PBS) that contained 0.3 M NaCl, 0.3% Triton X-100 and 5% normal donkey serum. Following immunoreaction, sections were rinsed for 5 minutes with PBS double salt prior to incubation in Hoechst (1:1000 in PBS) for 15 minutes, then rinsed for 5 minutes with PBS double salt. Sections were mounted in anti-fade medium and stored at −20°C.

Sections were scanned with a Zeiss 710 LSM confocal microscope (Carl Zeiss Microscopy GmbH, Germany) (Argon multi-line, 405 nm diode, 561nm solid state), using 20x objectives (NA 1.2), at 1 Airy Unit. 2 µm z-step images of the whole thickness of the section were obtained.

### Measurement of neuronal diameter

For the analysis of neuronal diameter, we examined transverse sections from the DRG of each animal. We used Neurolucida (version 2021.1.3, MBF Bioscience, USA) to measure neuronal area. To minimise selection bias, images were overlayed with a 160 x160 μm grid and the neuron closer to the upper left corner of each grid square was selected for analysis. Only NeuN-647 positive neurons with visible margins and nuclei included in the section were measured. To ensure that a total of 100 neurons were analysed per section, grid squares sizes were adjusted according to the total area of DRG section. To calculate neuronal diameter, data were transferred to Microsoft Excel (Microsoft Corporation, USA) and subsequently analysed in R (version 4.3.1) for graphical representation.

### Measurement of non-neuronal cell : neuronal cell ratio

For the analysis of the non-neuronal to neuronal cell ratio, the central focal plane of the immunohistochemical scans was selected and used for counting. Analysis was performed in Qupath (version 0.6.0,[5]). Neuronal nuclei were identified as Hoechst-positive and NeuN-647-positive, whereas non-neuronal nuclei were identified as Hoechst-positive and NeuN-647-negative.

### RNA Quality Assessment

10 µm thick sections of DRG tissue, were collected using a cryostat (Leica CM1950, Leica Microsystems, Germany), homogenized and processed for RNA extraction according to the standard RNeasy Micro-Qiagen protocol for RNA extraction from animal cells (Qiagen, Germany). RNA quality was assessed with RNA Integrity Number measured by Agilent 2100 Bioanalyzer (Agilent Technologies, USA).

### Cell Dissociation

DRG dissociation followed a described protocol with modifications[23]. Prior to dissociation all buffers and tubes were pre-chilled on ice. Briefly, DRGs were processed separately for each dog, such that each cell suspension corresponded to one individual animal (2-3 DRGs per dog). DRGs were placed in a petri dish on ice with 1mL of twin-salt-tris (TST) buffer. Surrounding roots and connective tissue were excised. While keeping the petri dish on ice, DRGs were minced with McPherson Vannas Scissors 45°angled (Fine Science Tools GmbH, Germany) and then Noyes Spring Scissors (Fine Science Tools GmbH, Germany) for 30 seconds.

To assess neuronal cell integrity and tissue dissociation, the minced DRGs were observed under a microscope. Mincing was repeated for the minimum time required to achieve full DRG dissociation. Following this, dissociation continued with fired-polished pipettes of decreasing diameter until a single-cell suspension containing neuronal cells was obtained. The resulting cell suspension was filtered through a 150 µm strainer (pluriSelect Life Science, Germany) into a 15 ml Falcon tube. Remaining cells in the petri dish were collected with an additional 1mL of TST and passed through the same filter, followed by 3mL of salt-tris buffer (ST). The filtered suspension was centrifuged at 4°C (500 x g, 5min).

The resulting pellet from each donor was resuspended in PBS containing 2% Bovine Serum Albumin (BSA) and incubated for 30 minutes at 4°C with conjugated NeuN-647 (ab190565, Abcam, UK) added at a final dilution of 1:500. Following incubation, samples were rinsed twice in PBS containing 2% BSA at 4°C. The final pellet was resuspended in PBS containing 0.04% BSA. To assess cell integrity, 10µl was taken and mixed with SytoxGreen (Catalog N° S7020, Thermo Fisher Scientific, USA, 1:1000) and visualised on an epifluorescence microscope prior to FACS.

### Cell Sorting

Prior to sorting, a colorimetric plate calibration was used as previously described[64]. Sorting was performed using a 130µm nozzle and Eppendorf twin.tec® PCR Plate 384 LoBind plates (Eppendorf, UK). Neuronal cells (Sytox Green-positive/ NeuN-647-positive) were sorted using a BD FACSAria IIU Flow Cytometer (BD Biosciences, UK) equipped with 4 laser (405, 488, 561, 633nm) and a 130 µm nozzle. Sytox Green emissions were detected using a 530/30BP, 502LP, and anti-NeuN-APC emissions using a 670/14 BP filter. The two-colour flow cytometric analysis was performed using BD FACSDiva Software (version 8.0.1, BD Biosciences, UK) on Microsoft Windows 7 operating system. At the start of each sorting session, unstained samples were used to verify gating.

Cells were sorted into 384 well plates containing FLASH-seq lysis buffer[32]. Each plate was sex-balanced during sorting, such as that half of the plate contained neuronal cells from a male, and the other half, cells from a female donor. Plates were centrifuged prior to sorting (500 x g, 1 minute) and were maintained at 4°C throughout FACS. Following sorting, plates were sealed, centrifuged (500 x g, 1 minute) and snap-frozen on dry ice and stored at −80°C until further processing.

### Sequencing & Alignment

The trial plate was processed and sequenced independently. The four final plates were processed in parallel. Single-cell RNA-seq libraries were generated using the FLASH-seq protocol with minor modifications[32].

Following cells sorting directly into 384-well plates containing lysis buffer supplied with dNTPs, SMART-dT30VN primer, dCTP, and FlashSeq-Template Switching Oligo (FS-TSO), the plates were snap-frozen and stored at −80°C until further processing. Reverse transcription polymerase chain reaction (RT-PCR) was performed by adding 4 µL of RT-PCR mix containing Superscript IV (Thermo Fisher Scientific, USA) to each well using the I.DOT Non Contact dispenser (Dispendix, Germany). The RT reaction was conducted at 50°C for 60 minutes followed by a pre-amplification PCR of 21 cycles. Amplified products were purified using the C.WASH noncontact centrifugal plate washer (CYTENA, Germany) for the beads clean up step.

To normalize the libraries, nuclease-free water was added to achieve a final average concentration of 200 pg/µL. Subsequently, 0.5 µL cDNA underwent tagmentation and enrichment using the Tagmentase (Tn5 transposase) – unloaded (Diagenode). Quantification of the final libraries was performed using the Qubit HS dsDNA assay and the fragment size distribution was measured using HS D1000 assay on an Agilent TapeStation 4200 system.

Sequencing was performed in paired-end mode (PE75) on a NovaSeq X Plus systems (Illumina, USA) using NovaSeq X Series 10B Reagent Kit chemistry. Raw sequencing data were demultiplexed using bcl2fastq2 v2.20 and aligned against C. lupus reference genome (Ensembl 108) using Kallisto v0.48.0.

### Analysis of single-cell RNA seq data

R (version 4.3.1) and Seurat (version 5.4.0) were used for the single-cell RNA-seq analysis. One Seurat object was generated per plate (five objects in total).

### Preliminary analyses to determine QC thresholds and batch correction strategy

Initial inspection of the data revealed differences in the number of detected unique genes per cell (nFeature_RNA) between sexes which drove significant cluster differences in un-batch corrected UMAP plots. To determine whether these differences reflected technical artifact or differences in sorted cell composition (i.e neuronal vs non-neuronal cells), datasets were first analysed separately by sex and no quality control thresholds (i.e. nFeature_RNA or nCount_RNA) were used.

Within each sex, layers were split by individual, to mitigate inter-individual batch effect. Data were first normalized (NormalizeData), highly variable features were identified (Find Variables Features), scaled (ScaleData), and subjected to Principal Component Analysis (RunPCA). To correct for dog-specific batch effects, datasets within each sex were integrated using canonical correlation analysis (CCA) (Integrate Layers, method=CCA Integration, origi.reduction=’pca’, new.reduction = “integrated.cca”, assay = “RNA”, verbose = FALSE). Clustering was performed using FindNeighbors(reduction = “integrated.cca”, dims = 1:30), FindClusters(resolution = 0.5, cluster.name = “cca_clusters”), RunUMAP(reduction = “integrated.cca”, dims = 1:30, reduction.name = “umap.cca”).

To compare cell-type composition between sexes, clusters were annotated into broad cellular classes using a marker gene list formed by canonical markers to distinguish neurons large and small diameter and non-neuronal (GFAP-positive) cells (marker_map <- list(Non_neurons =“GFAP”, Neurons= c(“UCHL1”, “TUBB3”), Large= c(“NEFH”,“NTRK2”, “PIEZO2”,“NTRK2”, “PIEZO2”,“NTRK3”,“PVALB”), Small=c(“INA”,“SCN10A”, “PRDM12”, “NTRK1”, “TRPV1”, “SST”, “CALCA”, “MRGPRD”, “VGLUT3”, “TRPM8”)). Comparison of annotated datasets showed similar proportions of neuronal vs non-neuronal cell populations between sexes, supporting nFeature_RNA differences were not caused by cell-composition and supporting subsequent preliminary joint analysis. A non-defined cluster characterised by uniformly low gene detection was labelled ‘Low’ and excluded.

Male and female datasets were then merged into a single Seurat object. Inspection of the merged dataset identified an additional cluster characterised by reduced nFeature_RNA and increased mitochondrial transcript abundance. All cells in this undefined additional ‘low’ cluster’ were removed using a nFeature_RNA threshold of 1250, and this was carried forward for final analysis.

### Final processing, integration and clustering

The filtered dataset (subset(subset = nFeature_RNA >= 1250)) was re-normalized (NormalizeData), and 3000 variable features were selected (Find Variables Features, nfeatures = 3000). Data were scaled (ScaleData) and subjected to Principal Component Analysis (RunPCA). Then layers were integrated using CCA, using dog as the integration variable (IntegrateLayers(method = CCAIntegration, orig.reduction = “pca”, new.reduction = “integrated.cca”, verbose = FALSE, k.weight=72); FindNeighbors(reduction = “integrated.cca”, dims = 1:30)); followed by final clustering (FindClusters(resolution = 1.5, cluster.name = “cca_clusters”)) and dimensionality reduction RunUMAP(reduction = “integrated.cca”, dims = 1:30, reduction.name = “umap.cca”).

Violin plots and FeaturePlot UMAPs were generated in Seurat to illustrate the distribution of selected genes across clusters. Unbiased marker genes for each cluster were identified using the Wilcoxon rank-sum test (FindMarkers) and filtered for avg_log2FC > 1 and adjusted p value < 0.05; the top 5 markers per cluster were selected based on adjusted p value and log2 fold change. Myelination index for each cluster was generated based on an aggregated expression score (AddModuleScore) of myelin associated genes (NEFH, NEFM, SCN8A, NFASC, KCNQ2, KCNQ3, KCNA1, CNTN1).

### Cross species Analysis

Mouse[68] and human[86] single cell/nucleus datasets were integrated with the canine dataset to enable comparisons across species. Prior to integration, all genes were converted to human 1:1 orthologues using the orthogene package (orthogene::convert_orthologs(method = “gprofiler”, drop_nonorths = TRUE, non121_strategy = “drop_both_species”)). New Seurat objects were then reconstructed using the filtered matrices while preserving the original metadata. This ensured that all downstream integration steps were performed on a common, human-symbol gene space across dog, human, and mouse. Orthologue-mapped dog, human, and mouse objects were then merged into a single Seurat object. Standard preprocessing steps were applied: log-normalisation, identification of 3,000 variable features, and scaling as described above.

For cross-species alignment, datasets were integrated using Seurat’s CCA-based layer integration (IntegrateLayers(method = CCAIntegration, orig.reduction = “pca”, new.reduction = “integrated.cca”)). Nearest-neighbour graphs were constructed using the integrated CCA reduction (FindNeighbors(dims = 1:30)), clusters were identified at resolution 1.2 (FindClusters(resolution = 1.2)), and UMAP embeddings were generated from the integrated space (RunUMAP(dims = 1:30)). Species-split and merged UMAPs were used to visualise cluster alignment and assess cross-species correspondence. These integrated UMAPs were used to assess cross-species correspondence among clusters and to confirm the presence of conserved neuronal types across dog, human and mouse. After integration, clustering was re-annotated into a simplified unified cross-species set, following the rationale outlined in the Results section. This approach allowed direct comparison of homologous sensory neuron classes.

To characterise molecular features that were shared or divergent across species, we performed cluster-wise differential expression analyses on the orthologue-mapped, integrated dataset. To focus analyses on physiologically and pharmacologically relevant gene families, we additionally generated curated gene lists representing ion channels, GPCRs, synaptic proteins, transcription factors, and neuropeptides (sourced from published datasets[16,33,74,77]). These lists were harmonised and used to define a focus gene set.

Prior to marker testing, we applied cross-species detectability filters to minimise artefacts arising from sparse counts or uneven species representation. For each gene, we computed the proportion of cells with detectable expression (>0 in the “data” slot) within dog, human, and mouse subsets. Genes were retained only if they were detected in at least 2% of cells in *each* species overall, and were detected in at least one species outside the focal cluster (>2% detection, i.e. globally detectable somewhere to reduce the potential for sparse potentially artifactual expression dominating results). Conserved markers were identified within each unified cross-species cluster using Seurat’s FindConservedMarkers (FindConservedMarkers, test.use = “wilcox”, only.pos = TRUE). Genes were required to have a positive log2 fold-change ≥ 0.25, expression in ≥ 10% of cells in the focal cluster, and species-wise adjusted P < 0.05. Conserved genes were ranked according to the maximum adjusted P-value across the three species, and the top 20 markers per cluster were visualised using dot plots with species as the grouping variable (DotPlot). Divergent, species-biased markers were identified using three pairwise contrasts within each cluster (Dog vs Human, Dog vs Mouse, Human vs Mouse) using FindMarkers (FindMarkers, logfc.threshold = 0.25, min.pct = 0.10, test.use = “wilcox”). Genes meeting an adjusted P-value threshold < 0.05 in any pairwise test were considered divergent for that species pair. For visualisation, the most strongly differentially expressed genes (top 20 by absolute log2 fold-change) were summarised in dot plots for each contrast. To quantify convergence and divergence across species, significant gene sets were partitioned into intersection categories (Human-only, Dog-only, Mouse-only, Human–Dog, Human–Mouse, Dog–Mouse, and Human–Dog–Mouse). Area-scaled Euler diagrams were generated using the eulerr package, based on the species-specific significance sets after applying all statistical and detectability filters. These diagrams represent the strict convergence of genes that are significantly enriched in the same cluster across all three species, which is more restrictive than the ranked top-20 conserved lists used for visualisation.

### Expression of Domestication selected genes in canine DRG

To test whether domestication-associated genes were non-randomly distributed across canine sensory neuron subtypes, we performed an enrichment analysis using a published list of positively selected genes (PSGs) identified in genome-wide domestication scans of dogs[89]. All analyses used PSGs that were detectably expressed within the dataset.

Genes expressed in at least three cells across the full cDRG dataset defined the background set (sum of non-zero counts ≥ 3). Differentially expressed genes (DEGs) were identified for each transcriptional cluster using Seurat’s FindMarkers (only.pos = TRUE, min.pct = 0.1, logfc.threshold = 0.25), and DEGs with p_val < 0.001 were retained. For each cluster, we tested whether PSGs were over-represented among DEGs using the upper-tail hypergeometric distribution. Here, X denotes the number of PSGs obtained when sampling n genes at random from the background set, allowing us to evaluate whether the observed overlap (k) exceeds that expected by chance. The enrichment probability was:

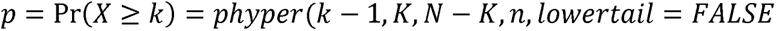

Where **N** = total number of background genes, **K** = number of background genes that were PSGs, **n** = number of DEGs for a given cluster, **k** = number of PSGs among those DEGs. P-values were corrected for multiple testing across all 13 clusters using the Benjamini–Hochberg false discovery rate (reported as *q*). This provided cluster-wise inference while controlling for global multiplicity.

To validate enrichments against a data-driven null, we performed 1,000 permutation tests per cluster. At each iteration: **n** genes were sampled uniformly at random from the background (preserving the observed cluster-specific DEG set size), the number of PSGs in the random sample was counted, and this distribution of random overlaps was compared to **k**, the observed overlap. The permutation p-value was computed as:

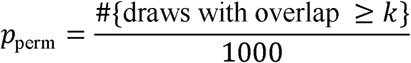

Clusters with 0/1000 permutations meeting or exceeding the observed value were reported as *< 1.00e−3*.

### Integration with canine 10x data

Data from FLASH-seq experiments was integrated with independently generated 10x Chromium canine DRG data to generate a high transcriptional coverage atlas. For methods related to this, please see the companion paper[41].

### Fluorescent in situ hybridization

Lumbar DRGs from Beagles were obtained as described above. Surrounding nerves and connective tissue were removed. A DRG from each dog was drop-fixed in Paraformaldehyde 4% for 13-16 hours, embedded in OCT and sectioned at 16 μm thickness using a cryostat (Leica CM1950, Leica Microsystems, Germany). Non-sequential sections were mounted onto Superfrost plus glass slides (Epredia, US), with one slide prepared per animal.

Fluorescent *in situ* hybridization was performed using HCR™ Gold RNA-FISH kit (Molecular Instruments, USA) according to the manufacturer’s protocol for slide-mounted samples. Slides were sequentially dehydrated in 50%, 70% and 100% ethanol (5, 5, and 10 minutes, respectively) at room temperature, air dried and each section group encircled using a hydrophobic pen (Hydrophobic Barrier Pen, Invitrogen). Slides were then placed in a warm humidified chamber and incubated in Hifi Probe Hybridization Buffer at 37°C for 10 minutes.

All probe sets were designed against dog (*Canis lupus familiaris*) mRNA sequences (Table S1). Probe sets were diluted 1:50 in Hifi Probe Hybridization buffer and applied to sections for overnight incubation at 37°C. Channel allocation varied between experiments (Table S1). Following hybridization, slides were rinsed four times for 15 minutes each in 1x HiFi Probe Wash Buffer at 37°C. Sections were then incubated in HCR Gold Amplifier Buffer at room temperature for 30 minutes prior to hairpin application. Corresponding hairpin amplifiers (Table S1) were snap-cooled, diluted 1:50 in HCR Gold Amplifier Buffer an applied to the sections for overnight incubation at room temperature in the humidified chamber in the dark. Following amplification, slides were washed four times for 15 minutes in 1x HCR Gold Amplifier Wash Buffer. Nuclei were counterstained with DAPI (1:1000) in 5x Saline Sodium Citrate (Invitrogen, Cat n° 15557-044, UK) for 1 minute. Slides were rinsed, air dried, and cover-slipped with mounting medium. Slides were stored at −20°C until imaging.

DRG sections were imaged on a Zeiss LSM 900 confocal microscope (Carl Zeiss Microscopy GmbH, Germany) (405, 488, 561, 640 nm diode) using a 40x magnification objective (NA 1.3) in Airyscan Mode. Laser channels selected to scan sections were chosen according to the Alexa Fluor channels selected for each probe and for DAPI, in addition, 488-nanometers channel was used in all scans to detect autofluorescence arising from lipofuscin within neuronal cell bodies, as it mimics true *in situ* signal[67].

One focal plane was acquired for each DRG section. To ensure sufficient sampling (at least 100 neurons per dog) at least three sections were imaged per dog. The sections imaged were selected at random, but preference was given to sections free of sectioning artifact.

### Analysis of ISH images

Airyscan raw image files were processed and stitched using Zen software (Carl Zeiss Microscopy GmbH, Germany) and individual images were generated for each section. Images were opened in QuPath (version 0.6.0). For all analysis, only neuronal cells with visible nuclei in the acquired focal plane, identified by DAPI staining, were included. Probe signal was assessed manually, using the 488 nm autofluorescence channel to identify and exclude lipofuscin-associated signal from scoring. All scoring was performed by a single observer per experiment.

Neuronal categorisation followed similar analytical workflows for both probe combinations, with channel specific modifications as described below. For analyses based on IL31RA expression, IL31RA-positive neurons were annotated initially, after which IL31RA annotations were masked to avoid bias during subsequent GFRA2 identification. Co-expression was then determined by overlay of the two independent annotations.

For KIT and SSTR2 probes, heterogeneity in signal intensity and puncta density was observed across neuronal cell bodies. Neurons were therefore classified into three categories: negative (no signal above background), positive (+; sparse puncta distributed across the cytoplasm without homogenous clustering), or strongly positive (++; dense, clustered puncta filling the cytoplasm with high signal intensity). Soma area was measured by manually delineating the soma boundaries, neuronal diameter was calculated assuming neuronal circular geometry using the equation 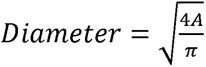.

Neurons were categorised in small (< 40μm), medium (40-60μm) and large (>60μm). For both analyses, neuronal measurements and classification data were transferred to Microsoft Excel (Microsoft Corporation, USA) and subsequently analysed in R (version 4.3.1) for statistical analysis and graphical representation.

### Reanalysis of Xenium in situ data

Publicly available Xenium data from Yu et al.[86] was reanalysed in Xenium explorer software (version 4.1.1, 10x Genomics, USA). DRG sections were viewed with the DAPI channel visible. Neurons expressing >5 cytoplasmic transcripts of either IL31RA or GFRA2 were marked using the annotations feature as GFRA2+ or IL31RA+. Neurons of each class were annotated separately without viewing the channel for the other transcript. Annotations were exported and colocalization calculated using custom R scripts.

## Results

### Assessment of neuronal size and neuronal versus non-neuronal cell ratio in cDRG

To inform our neuronal isolation and sequencing methodologies we first sought to define the neuronal size distribution and ratio of neurons to non-neuronal cells in cDRG. For these initial experiments we used tissue sections from 4 dogs of mixed breeds (German Shepherd, Greyhound, XL Bully & Border Collie) which were reacted with antibodies against NeuN and counterstained with DAPI. We observed that NeuN labelled all neuronal profiles (as recognised using DAPI staining by size, profile and the presence of a large pale nucleus compared to non-neuronal cells, Fig 1). Using a stereological technique, we measured the size of 100 neurons per animal and found that neuronal diameter ranged from 21 μm to 81 μm (Fig 1 & Table S2). When compared to sizes of DRG neurons across species [69,85], cDRG neurons appear to be of a size more akin to human than rodents. We also observed that neuronal nuclei were greatly outnumbered by non-neuronal nuclei. We quantified this in single optical sections from the same confocal images as used above, with this method chosen to allow comparisons with recently available data from human DRG (hDRG)[15]. We manually counted an average of 15,549 (range 14,626-16,045) non-neuronal nuclei per animal versus 224 (194-265) neuronal nuclei. The ratio of neuronal to non-neuronal nuclei across the whole DRG section, was calculated as 1:65 (range 1:56-1:75, Fig S1). Overall, Neuronal nuclei comprised approximately 1.6% of all nuclei, closely matching reported proportions in human DRG[15]. The abundance of non-neuronal cells led us to develop a method to enrich neuronal cells for sequencing using fluorescence activated cell sorting (FACS).

**Fig 1.**
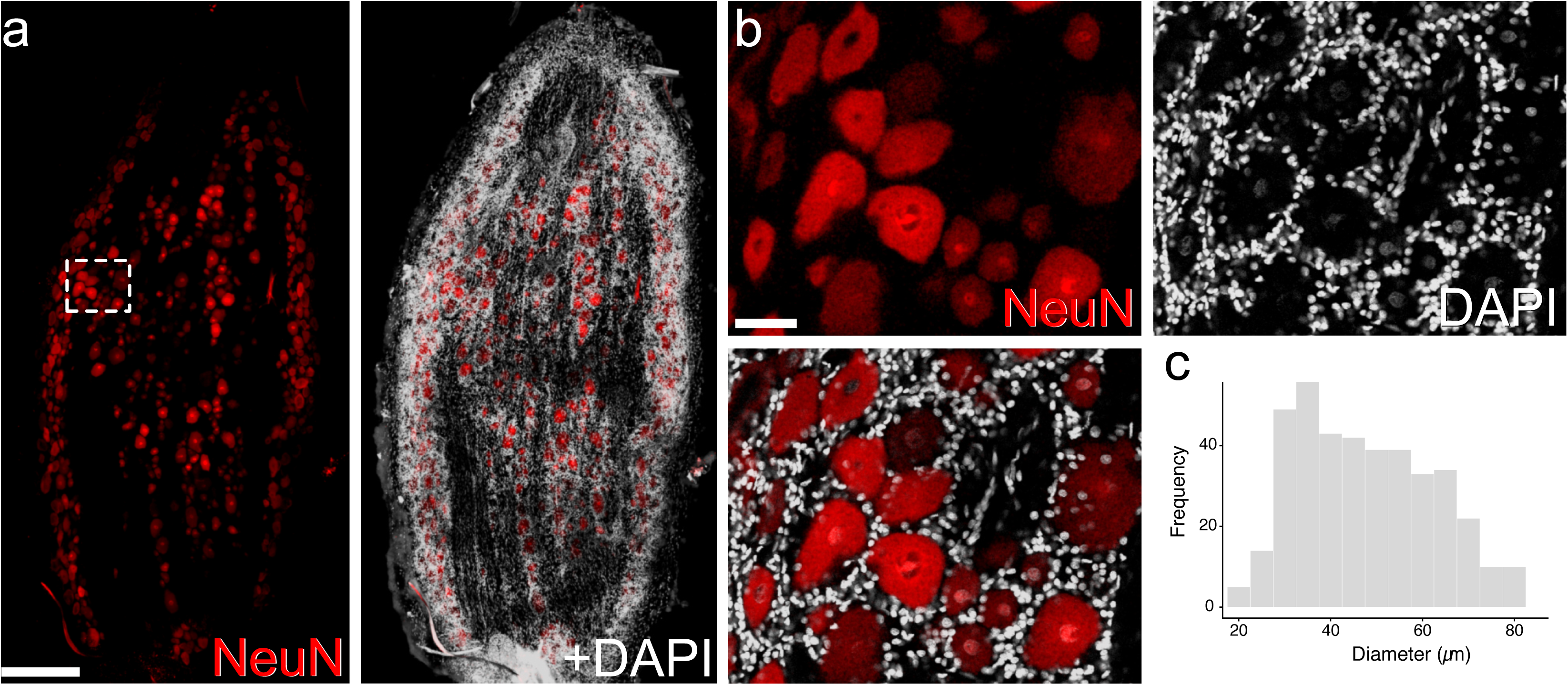
Neuronal size and abundance of non-neuronal cells in the canine DRG. **(a)** a confocal image of canine DRG stained for NeuN (neurons) and DAPI (all nuclei). **(b)** Higher magnification views are shown in panels. Large and small neurons are visible surrounded by many non-neuronal cells. Low resolution image is a maximum projection of 7 optical sections acquired at 2µm z-resolution while the inset is a single optical section, scale bars are 400 µm and 50 µm, respectively. (**c)** Histogram showing the size profile of neurons counted across sections from 4 animals, n = 100 neurons per animal.

### Dissociation of cDRG and FACS purification of neurons for scRNA-seq

Lumbar ganglia for all FACS and sequencing experiments were acquired post-mortem from 10 laboratory beagles (6 males and 4 females, 9-24 months old) and flash frozen. These dogs were control (no drug) animals in unrelated toxicological studies, and were healthy with no known painful conditions. To ensure high-quality RNA, we verified RNA integrity number (RIN) was greater than 7.5 for each donor (Table S3/Fig S2a). In initial experiments, the ganglia’s robust nature and connective tissue presented a challenge to dissociation. However, we found that using a dissociation protocol based on mincing and triturating tissue in Tween-based salt solution[32] could successfully dissociate the ganglia. Following dissociation, we performed NeuN staining of the resultant suspension and consistently observed that whole neuronal cells, with recognisable cytoplasm and nuclei, were present alongside non-neuronal nuclei (Fig S2c). We purified these neuronal cells via FACS (Fig S2b&c), sorting cells directly into individual wells of 384-well plates, which were subsequently submitted for transcriptomic analysis using the FLASH-seq method [32].

Prior to performing the definitive RNA-seq experiment, we ran a small pilot experiment to optimise FACS strategy prior to further experiments. In this trial library, neurons represented 89 % of cells passing QC with a small cluster of non-neuronal cells expressing GFAP evident (Fig S2d). We conducted a principal component analysis using FACS gating parameters to determine if cells in the GFAP+ cluster could be excluded via gating. However, GFAP+ cells were not distinguishable on this basis (Fig S2e). A moderate number of wells failed to generate data, likely due to plate sorting inaccuracies, resulting in 215 cells passing QC in a 384-well plate. Despite the lower-than-expected yield, these findings demonstrate the feasibility to significantly enrich for neuronal cells prior to sequencing using this FACS protocol.

### Clusters of canine DRG neurons and marker genes

To generate our final dataset, we sorted an additional 4 plates of cDRG cells and submitted these for sequencing. We used lumbar DRGs from 8 donor beagle dogs of both sexes. Plates were filled over two sessions on separate days, using a balanced design where each plate contained cells from a male and female animal. Following quality control, and removal of a GFAP+ cluster of 93 cells, 934 cells were carried through to the final dataset with a mean unique genes per cell of 8,923 (Fig S3a). We examined QC metrics across plate, sex, dog and session and observed a pattern where cells from female animals were consistently of lower quality (Fig S3b&c), perhaps due to fragmentation of cytoplasm. However, Seurat Canonical Correlation Analysis (CCA) integration with dog as the batch layer resulted in the same distribution of clusters across sex and plate (Fig S3d&e). In order to perform CCA we removed cells from one animal (male) from the dataset due to this animal having a number of QC-passing cells (21) that was too low for the integration algorithm, resulting in a final dataset of 913 cells.

Across the clusters, the cells were deemed to be excitatory neurons due to the expression of the neuronal markers UCHL1 and TUBB3, and the glutamate transporter SLC17A6 (Fig 2a). Genes known to be expressed across broad populations of sensory neurons, such as the Nav1.8 sodium channel (SCN10A), the heat=sensitive ion channel TRPV1, the μ=opioid receptor (OPRM1), the neurotrophin receptors NTRK1 and NTRK3, and the mechanosensitive ion channel PIEZO2, were detected across multiple clusters (Fig 2b). We selected a cluster resolution of 1.5, resulting in 12 clusters (d0-d11) to annotate (Fig 2c & S4), we then elected to additionally subdivide the small clusters d5 & d8 as these appeared to include biologically meaningful subtypes of myelinated neurons (Fig 2d). Using known markers for populations of sensory neurons[11,42] (Fig 2e & S5), and unbiased differentially expressed genes (DEGs) (Fig 2b) we annotated the resultant clusters as follows (Fig 2f).

**Fig 2.**
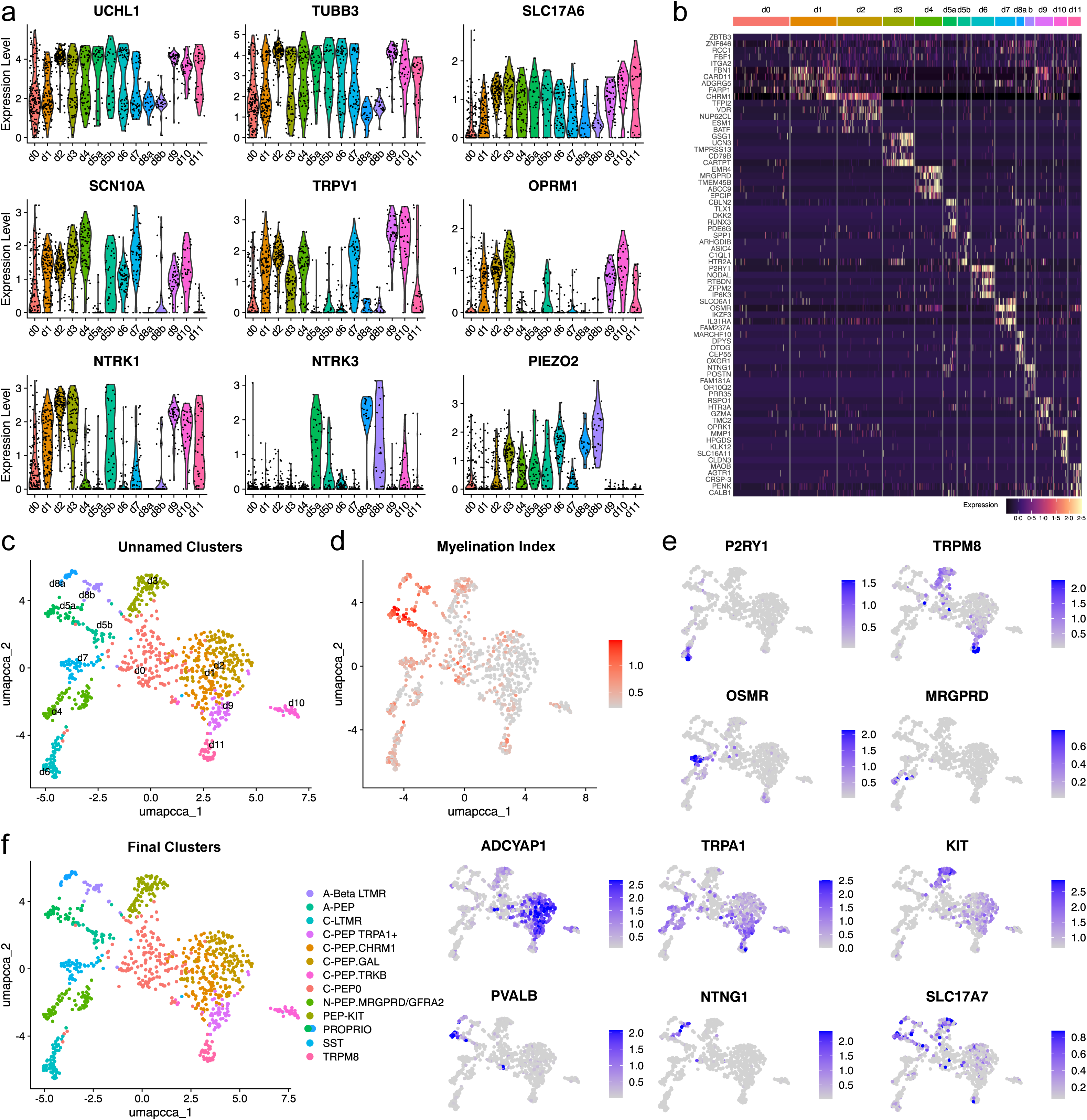
Transcriptomic definition and annotation of canine DRG neuronal subtypes. **(a)** Violin plots showing expression of broad neuronal markers (*UCHL1*, *TUBB3*, *SLC17A6*) and core somatosensory genes (*SCN10A*, *TRPV1*, *NTRK1*, *NTRK3*, *OPRM1*, *PIEZO2*) across all clusters. **(b)** Heatmap of unbiased differentially expressed genes (DEGs) distinguishing the initial unsupervised clusters. **(c)** UMAP projection of FLASH-seq profiles showing clustering of 913 cDRG neurons into transcriptionally distinct groups (d0–d11). **(d)** Myelination index for each cluster based on an aggregated expression score of myelin-associated genes (*NEFH*, *NEFM*, *SCN8A*, *NFASC*, *KCNQ2*, *KCNQ3*, *KCNA1*, *CNTN1*), highlighting putative myelinated versus unmyelinated populations. **(e)** UMAP feature plots illustrating selected cluster-specific marker genes used for manual annotation. **(f)** Final annotated UMAP showing the resolved sensory neuron subtypes, including C-fibre nociceptors (C-PEP0, C-PEP.TRPA1, C-PEP.GAL, C-PEP.CHRM1, C-PEP.TRKB), non-peptidergic nociceptors, SST/OSMR neurons, C-LTMRs, PEP-KIT neurons, proprioceptors, A-δ nociceptors and A-β LTMRs.

Firstly, a cluster of putative cold sensing neurons (d11) was identified by the expression of the temperature sensing ion channel TRPM8, with this cluster also expressing FOXP2 and SCN5A; both established markers of cold sensitive cells[11,42]. Interestingly, cells in this cluster also expressed high levels of PENK, encoding the opioid precursor proenkephalin. We also annotated a cluster (d6) as C-low threshold mechanoreceptors (C-LTMR) on the basis of expression of the marker genes CDH9, TAFA4 and P2YR1[25]. As is the case in humans[58,86], cells in this cluster did not express the canonical rodent marker genes SLC17A8 or TH[11].

Two clusters in our dataset resembled non-peptidergic nociceptors. We annotated the first of these (d7) as Somatostatin (SST) non-peptidergic neurons based on the expression of genes predominantly related to transducing itch inducing stimuli; including the cytokine receptors IL31RA and OSMR, the potassium channel KCNQ4, and the histamine receptors HRH1 and HRH2[1,20,83]. This cluster was also enriched for JAK1, a crucial and targetable component of the downstream effector pathway for cytokine mediated itch[12,57]. Cells in this cluster also expressed a number of unexpected genes including PIEZO1, NPYR2 and the cytokine receptor IL17RD. The second cluster (d4) annotated as non-peptidergic expressed classical cross-species markers of non-peptidergic neurons including MRGPRD, GFRA1, GFRA2 and LPAR3[11,42]. As has been reported in human DRG[69,76], these cells also express moderate levels of peptidergic marker genes such as CALCA and TRPV1, providing evidence that, as seen in other sequencing datasets, a binary division of nociceptors on the basis of peptide expression is not possible in the dog. We also observed selective expression of the ion channel TRPM6 in this population, a channel that is not commonly recognised as a specific feature of sensory neuron clusters in other species.

We identified a number of clusters (d0, d1, d2, d9 & d10) that corresponded to putative unmyelinated C-fibre nociceptors based on the expression of low levels of genes associated with myelination, plus expression of the markers TRPV1, CALCA, NTRK1, OPRM1 and ADCYAP1[11,42,79]. This set of clusters was annotated on the following basis. The d9 cluster expressed high levels of the TRPA1 ion channel, akin to a small cluster of C-fibre nociceptors described in humans[86], and was therefore labelled as C-PEP.TRPA1. Cells within this cluster also expressed low levels of the opioid receptor OPRK1, and the serotonin receptor HTR3A. We labelled the two adjacent clusters d1 and d2, based on unbiased marker gene detection (Fig 2b), as C-PEP.GAL and C-PEP.CHRM1. Cells in the C-PEP.GAL cluster expressed galanin and the previously recognised markers of subsets of peptidergic nociceptors TACR3 and PROKR2[11], while cells in the C-PEP.CHRM1 cluster expressed the muscarinic receptor gene CHRM1. We named cluster d10 as C-PEP.TRKB due to the expression of the neurotrophin receptor NTRK2, however no single markers linked this cluster to an established class of DRG neuron, and the cross-species analogue of this cluster remains unresolved, although overlap of CALCA and NTRK2 is seen in a single subtype in cross-species atlases (Calca+Dcn in [11]). Finally, we identified a number of cells (cluster d0) which were characterised by a lack of distinctive markers despite the presence of generalised neuronal markers. Previous human DRG datasets, generated using full-length SMART technologies have also identified similar clusters and termed these hPEP0[86]. We therefore designated this cluster as C-PEP0 in our annotation.

Cluster d3 was characterised by high levels of expression of the proto-oncogene KIT, and low levels of TRPM8 and TRPV1. A similar cluster of peptidergic nociceptors has recently been described in humans [86] and we annotated our d3 cluster as PEP-KIT to reflect this apparent similarity. The PEP-KIT cluster also expressed moderate levels of SLC17A7 and VSNL1, which are recognised as cross-species markers of myelinated afferents [42], however our gene-expression based myelination index did not indicate this cluster was heavily myelinated. As such it is currently unclear if this PEP-KIT cluster is myelinated in dogs.

Finally, we annotated the two clusters likely containing large, myelinated neurons, as suggested by the myelination index and expression of NTRK3. These cells appeared to be underrepresented in our dataset, perhaps due to the FACS sorting protocol biasing towards smaller cells. Using a higher cluster resolution than could be applied globally, we further subsetted these clusters into clusters d5a, d5b, d8a and d8b to identify different classes of myelinated afferents. Clusters d5a and d8a were annotated as proprioceptors due to the expression of parvalbumin (PVALB). Cluster d5b was identified as a myelinated putative A-delta nociceptor class due to expression of CALCA and SCN10A. This cluster also expressed the mechanosensitive ion channel PIEZO2 and the serotonin receptor HTR2A. Lastly, the d8b cluster was designated as a group of A-beta LTMRs due to the expression of high levels of PIEZO2, the axonal guidance molecule NTNG1[11] and the A-fibre marker KCNH5[86]. We also asked if FACS parameters could retrospectively be used to distinguish putative large neurons, as defined by the expression of NTRK3 (a gene selectively expressed in a subset of large neurons), from small neurons, as cell size is known to influence flow parameters. We found no robust FACS parameter-based distinction based on either principal component analysis of all FACS data, or pairwise examination of FACS gating measures (Fig S6).

### Cross-species integration reveals convergent and divergent subtype-specific gene expression

Next, we sought to compare cell-type specific gene expression across species via integrating our canine FLASH-seq cells with existing human and mouse single-cell datasets using 1:1 orthologues. We used Seurat CCA integration to generate cross-species UMAP plots including human cells sequenced using SMARTseq2 [86] and an adult mouse dataset with high cell numbers and hence transcriptional coverage [68]. There was significant overlap between clusters (Fig 3a), in particular between the two full-length technologies (human and canine). In order to make cross-species comparisons, we elected to reannotate these cross-species clusters using a simplified unified annotation with 10 clusters (Fig 3b & Fig 3c). Although the unified cross=species annotation is necessarily coarse, this approach reflects a deliberate trade=off: merging of clusters due to lower cell numbers than droplet-based studies versus exceptionally high molecular depth, thereby permitting robust cross-species comparisons across the annotated clusters.

**Fig 3.**
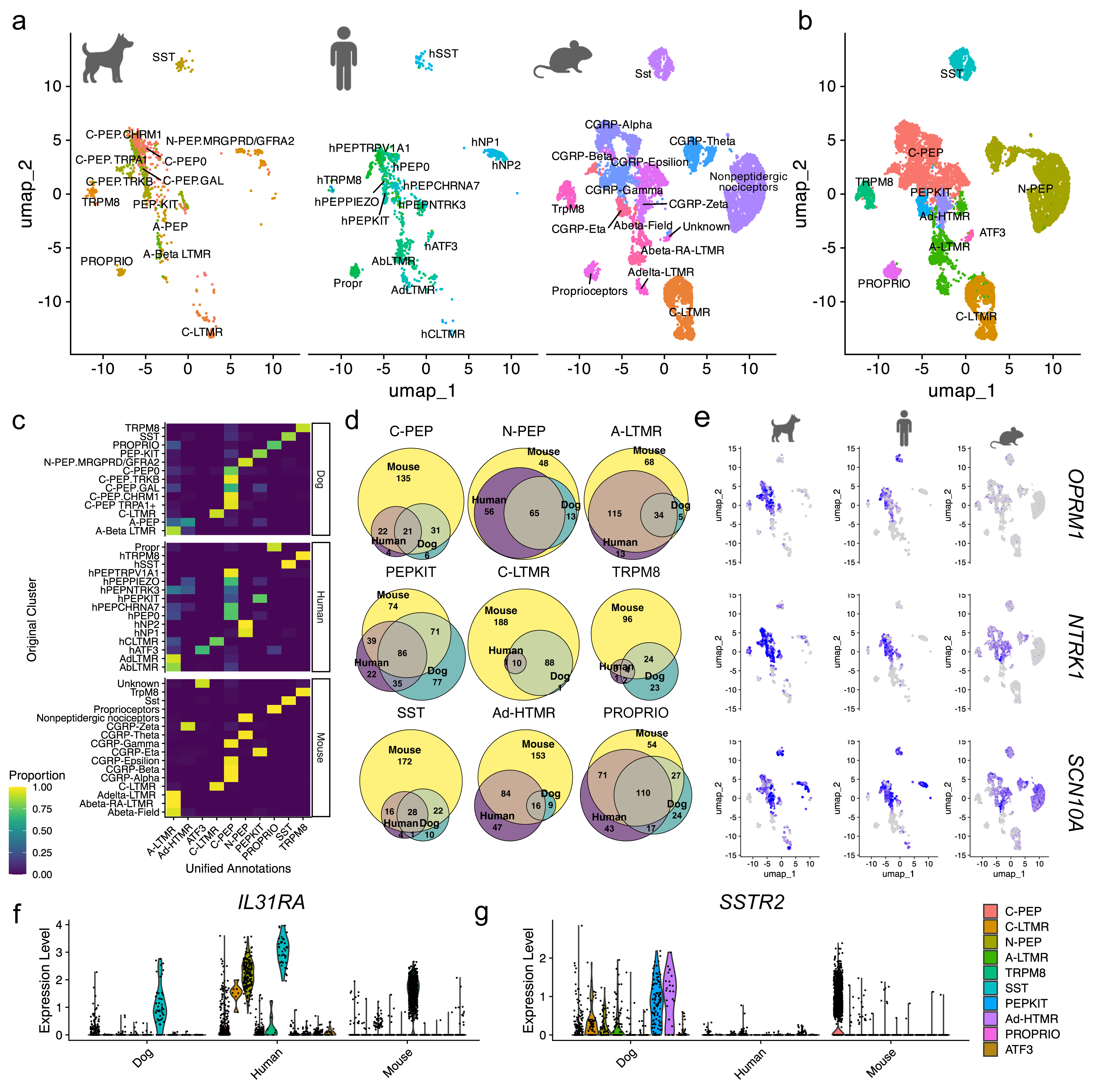
**(a)** UMAPs showing integrated dog, human, and mouse DRG datasets coloured by original study-specific cluster identities, illustrating cross-species alignment obtained using 1:1 orthologues and CCA integration. **(b)** Unified annotation of the integrated dataset collapsed into 10 consensus sensory neuron classes, enabling direct comparison of homologous subtypes across species. **(c)** Heatmaps showing how original dog, human, and mouse clusters reallocated into the unified consensus annotation following integration. **(d)** Summary of convergent and divergent DEGs across species within each consensus cluster, highlighting transcripts with conserved enrichment versus species-biased specialisation. **(e)** Examples of conserved pain- and itch-relevant genes that show strong cross-species alignment in cluster-specific expression patterns. **(f–g)** Cross-species expression patterns of *SSTR2* and *IL31RA* across neuronal subtypes, illustrating *IL31RA* expression restricted to canine SST/OSMR neurons and the broader *SSTR2* distribution in canine DRG, as quantified in subsequent ISH analyses.

To determine which molecular features of sensory neurons are shared across species, we first assessed convergent gene expression within each unified cluster. Using a list of physiologically and pharmacologically relevant gene families (ion channels, GPCRs and their ligands, synaptic proteins, transcription factors, and neuropeptides), we identified genes that were robustly detected in both dog and human datasets and were consistently enriched within the same cluster (Fig 3d). This analysis revealed a substantial set of highly conserved markers, many of which represent established therapeutic targets for pain and itch. Notably, transcripts encoding the μ□opioid receptor OPRM1, the neurotrophin receptor NTRK1, and the nociceptor□enriched sodium channel SCN10A showed strong and parallel enrichment across aligned dog–human clusters, indicating the conservation of core nociceptor signalling pathways (Fig 3e). Comparable patterns were also observed when integrating dog and mouse datasets (Fig S7).

We next examined divergent expression patterns, i.e. genes that showed consistent differences between dog and human within the same neuronal cluster. Because artefactual divergence can arise from sparse detection, cluster size differences, or platform□specific dropout, we incorporated minimal detection requirements before testing. Only genes with detectable expression in both species across the datasets were evaluated further. Despite this approach, it is likely that many of the divergent genes likely result from technical factors. However, on examination of the results (Figs S8 & S9) we were able to identify a number of subtype□restricted, species□biased transcripts with relevance to pain and itch. As detailed in subsequent sections, these include IL31RA, which in dogs is highly restricted to OSMR/SST neurons, and the somatostatin receptor, SSTR2, which shows strikingly broad expression across neuronal subtypes in the dog (Fig 3f & Fig 3g).

### In situ validation of divergent patterns of gene expression relevant to itch and pain therapeutics

We then proceeded to validate the apparent divergent IL31RA expression pattern between dogs and humans using *in situ* hybridisation (ISH). Our cross-species analysis indicated that IL31RA is restricted to the SST/OSMR cluster in canine DRG; in contrast to human DRG, where IL31RA is additionally expressed in non-peptidergic nociceptors and C-LTMRs (Fig 3f). To verify this, we performed ISH using probes against IL31RA alongside GFRA2; a conserved marker for C-LTMRs, and non-peptidergic nociceptors in both dogs and humans (Figs S10a & S10b). We used canine DRG tissue from 4 animals, of both sexes, and restricted our analysis to neurons expressing IL31RA or GFRA2 (Fig 4a). We counted a mean of 50 (range 36-67) IL31RA+ and 180 (115-260) GFRA2+ neurons per animal. No neurons co-expressed IL31RA and GFRA2. To determine if IL31RA expression is non-specific to SST neurons in human DRG, we reanalysed existing, publicly available, human *in situ* Xenium data [86]. Across 4 Xenium sections we counted 241 GFRA2+ and 266 IL31RA+ neurons. Of these, 133 cells co-expressed both genes with 50% of IL31RA+ neurons expressing GFRA2. (Fig S10c). Overall, these results confirm a restricted IL31RA expression pattern in dogs, in striking contrast with human DRG. This species difference has been previously suggested in a canine vs human DRG bulk sequencing dataset [38], our reanalysis of which shows IL31RA transcript abundance to be lower in canine DRG than in humans (Fig S10d). However, here we show that this difference relates to cell-type specific expression patterns rather than simply differences in absolute expression.

**Fig 4.**
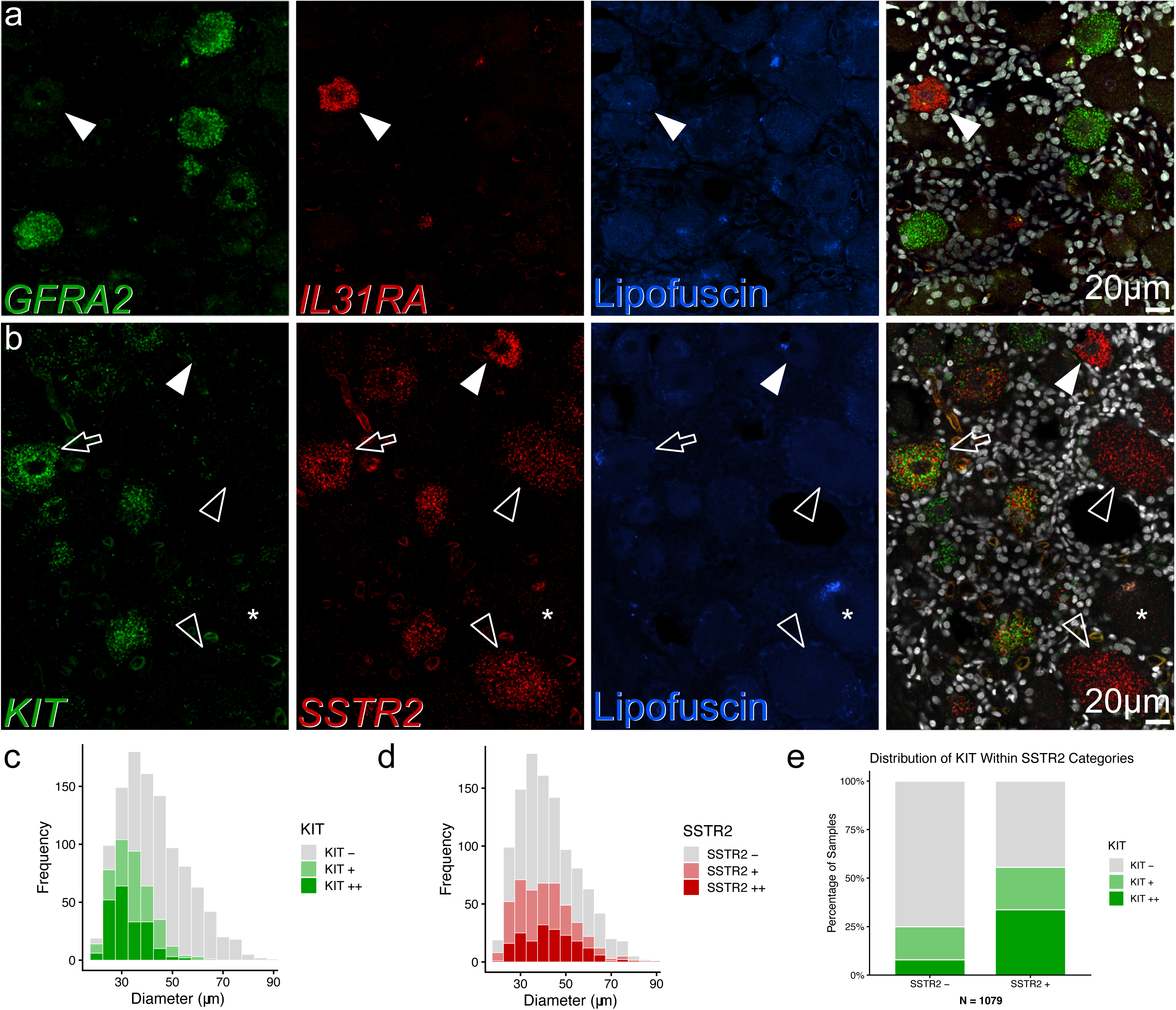
*In situ* validation of divergent *IL31RA* and *SSTR2* expression patterns in canine DRG. **(a)** HCR ISH detecting *GFRA2* and *IL31RA* transcripts in canine DRG tissue. Lipofuscin/autofluorescence is shown in blue. Filled arrowhead indicates a single *IL31RA*+ neuron. There is no detectable expression of *IL31RA* in adjacent *GFRA2*+ neurons, confirming the mutually exclusive expression of these markers in dogs. Images represent single confocal optical sections acquired at 40× magnification. Scale bar = 20□µm. **(b)** HCR ISH for *KIT* and *SSTR2*. A filled arrowhead indicates a small-diameter neuron expressing *SSTR2*, whereas open arrowheads denote medium-to-large neurons expressing *SSTR2*. *KIT* expression is seen in subsets of small to medium diameter neurons. Unfilled arrow shows a medium diameter *KIT+* neuron with high *SSTR2* expression. A large-diameter neuron lacking *SSTR2* expression is marked by a star. Images are single confocal optical sections acquired at 40× magnification. Scale bar = 20□µm. **(c)** Soma-diameter distribution of *KIT*□ neurons, demonstrating that *KIT* expression spans small- and medium-sized neuronal populations. **(d)** Soma-diameter distribution of *SSTR2*+neurons illustrating expansion of expression beyond the small-diameter range into medium-sized neurons. **(e)** Quantification of *KIT* expression within *SSTR2*□ and *SSTR2*□ neurons.

Interestingly, SSTR2 expression in dogs presents a contrasting pattern to that seen in rodents. In the mouse, SSTR2 is primarily detected in small-diameter C-fiber nociceptors and is indeed restricted such that it is a canonical sub-population marker[62]; whereas in dogs, SSTR2 is seen in small-diameter neuronal populations (C-LTMRs, non-peptidergic nociceptors) but is also highly expressed in PEP-KIT and A-delta nociceptors (small to medium-neuronal diameter). Expression is also seen in some putative A-LTMRs in our dataset. Therefore, we hypothesized that SSTR2 would be robustly expressed across small and medium diameter neurons in cDRG, with expression extending into some medium to large cells. We also hypothesized that a significant proportion of SSTR2+ neurons would express high levels of KIT and hence would belong to the PEP-KIT population. To investigate this, we again performed ISH in DRGs from 4 donors using SSTR2 and KIT probes (Fig 4b). We first measured the size of all neurons with nuclei present in the sections (n = 1079, Figs 4 c & d). Probe signal for SSTR2 and KIT was divided into 3 categories (negative, + and ++; see methods) and neurons were classified according to their soma diameter into small (< 40μm), medium (40-60μm) and large (>60μm). We identified a total of 457 neurons expressing SSTR2, of which 254 (58%) co-expressed KIT (Fig 4e). Based on diameter, SSTR2 was mostly seen in small to medium neurons although some expression was seen in large neurons (Fig 4d, Table S5); whereas, in contrast, KIT was primarily a marker of small to medium sized neurons (Fig 4c, Table S5). Overall, these results show that SSTR2 expression in dogs is seen in a wider neuronal population than in mice and this includes cells likely to be nociceptors but also some putative LTMRs based on size profiles.

### Gene expression in canine DRG neurons is likely influenced by domestication

It has been suggested that evolutionary pressures due to domestication have influenced gene expression in other areas of the canine nervous system such as the hippocampus [89]. DRG sensory neurons arise from the neural crest, an embryonic lineage proposed to be a recurrent target of domestication□related selection[60], and this raises the possibility that transcriptional specialisation within somatosensory neurons may also carry signatures of domestication. In order to establish whether this could also be the case, we tested whether putative positively selected genes (PSGs) from published genome-wide domestication scans [4,30,60,81,82] were non□randomly distributed, i.e., enriched, across cDRG transcriptional subtypes. Of all described PSGs, 573 were detectably expressed and were included in enrichment analyses.

Hypergeometric analysis revealed significant over□representation of PSGs in a subset of DRG neuronal clusters (Table 1). A separate 1,000□permutation analysis confirmed the same set of non-random enrichments of PSGs in clusters. The strongest enrichments were observed in peptidergic C-fibres, non-peptidergic C-fibres, C-LTMRs, Proprioceptors and SST neurons. The convergence of both statistical approaches supports a biologically meaningful overrepresentation of domestication□associated genes within specific DRG neuronal lineages and may demonstrate that canine domestication has preferentially impacted specific sensory neuron pathways.

**Table 1.** Enrichment of positively selected genes (PSGs) across canine DRG neuronal subtypes. For each transcriptional cluster, the number of PSGs overlapping the cluster-specific DEG set is shown together with enrichment significance assessed by hypergeometric testing (BH-adjusted across 13 clusters) and by 1000 permutation testing (values of 0/1000 reported as <1.00e−03). Selected PSGs list representative domestication-associated genes present in each cluster’s PSGvsDEG set.

| CLUSTER | PSG OVERLAP<br>(N) | HYPERGEOMETRIC<br>P-VAL ADJUSTED | PERMUTATIO<br>N P-VAL | SELECTED PSGS (EXEMPLARS) |
| --- | --- | --- | --- | --- |
| C-PEP.GAL | 207 | 1.06e-07 | <1.00e-03 | OPRM1, GNAQ, NGFR, TLX3, GPR139 |
| C-PEP.TRPA1 | 136 | 1.35e-05 | <1.00e-03 | OPRM1, FOXP2, GPR139, COL11A1, CACNB3 |
| N-PEP.MRGPRD/GFR<br>A2 | 50 | 3.13e-03 | 1.00e-03 | CISH, PLCE1, KCNC2, GRB10, GPR139 |
| C-LTMR | 50 | 3.23e-03 | <1.00e-03 | GAD2, FGF1, GRB10, CISH, CNTN5 |
| PROPRIO | 70 | 3.23e-03 | 3.00e-03 | CALB2, NTNG1, CNTN5, FGF1, MAPKAPK3 |
| SST | 35 | 6.37e-03 | 1.00e-03 | IL31RA, GPR139, PLCE1, CYP1B1, TIAM1 |
| C-PEP.TRKB | 61 | 1.22e-02 | 7.00e-03 | OPRM1, NGFR, GAD2, TCF4, GNAQ |
| C-PEP.CHRM1 | 24 | 4.28e-01 | 2.84e-01 | GPR139, SCN2A, COL11A1, NRXN3 |
| A-PEP | 9 | 4.47e-01 | 3.54e-01 | NGFR, SEMA3D, GALR1, ADCY8 |
| C-PEP0 | 36 | 4.47e-01 | 3.07e-01 | IKZF1, C5AR1, PITX1 |
| PEP-KIT | 100 | 4.47e-01 | 3.56e-01 | OPRM1, NRXN3, CACNA1A, LRRTM3 |
| TRPM8 | 12 | 7.75e-01 | 6.96e-01 | FOXP2, CALB2, NKAIN2 |
| A-BETA LTMR | 160 | 7.75e-01 | 7.76e-01 | NTNG1, GALR2, CNTN5, CACNA1A, ADM |

Furthermore, we selected example genes with roles in pain and itch (Table 1) from the full lists of enriched PSGs (Table S6) to demonstrate that many potentially selected genes are linked to neuronal signalling and sensory function. Of note, IL31RA has been previously documented as being influenced by domestication and its enrichment within specific cDRG neuronal subtypes suggests that evolutionary pressures may have influenced the regulatory context of IL31RA in dogs [60]. Overall, the over□representation of PSGs in cDRG transcriptional clusters suggests that selection associated with domestication may have systematically shaped pain, itch, and mechanosensory circuits in the dog thereby generating species specific transcriptional programs for sensory function.

### Canine atlas generation and label transfer from a large human dataset confirms cluster identities

Finally, we validated our single□cell clusters by integrating them with an independently generated single□nucleus canine DRG dataset produced by a collaborating group[41].This cross□platform integration combined our deep, full□length FLASH□seq profiles with a large□scale 10x Chromium snRNA□seq dataset. This substantially increased total cell numbers and expanded transcriptional coverage, while preserving the high□resolution molecular detail that distinguishes our dataset. The two canine datasets showed strong concordance, with clear alignment of major sensory neuron classes, robust cross□platform reproducibility of cluster identities, and improved representation of large□diameter neuronal populations that were relatively under□captured by FACS (Fig 5a). Together, these analyses enabled consensus cluster naming, finer□grained annotation of neuronal subtypes, and the construction of a unified canine DRG atlas that will serve as a valuable comparative resource for future sensory neurobiology. Full details of the atlas generation and benchmarking are provided in the companion paper[41].

**Fig 5.**
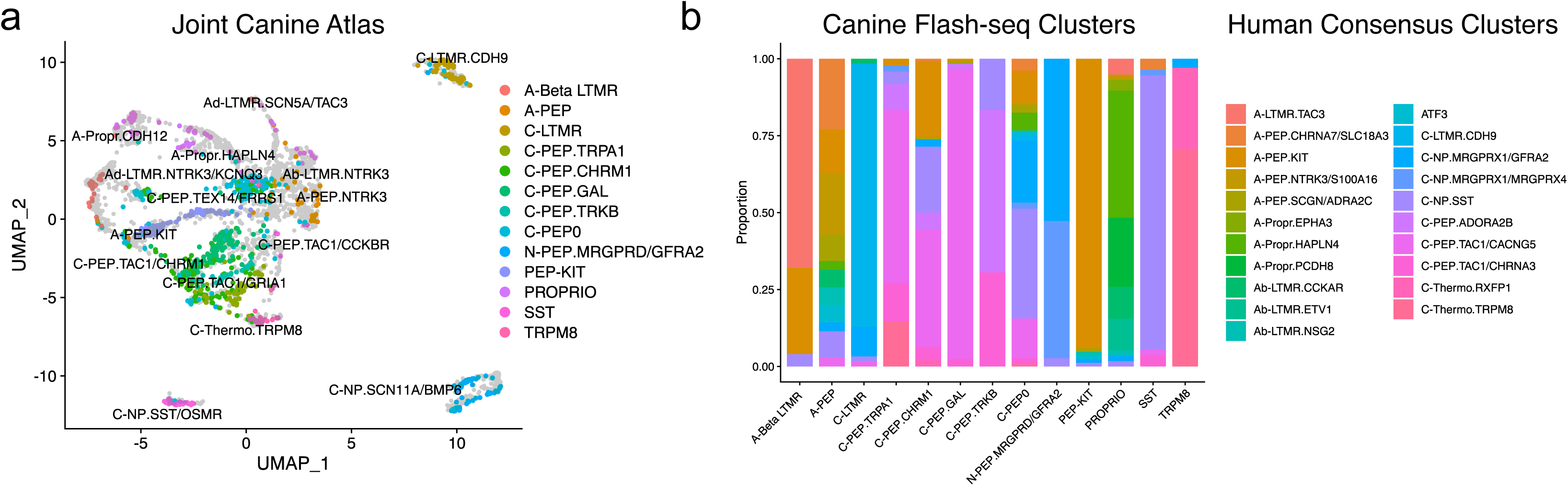
Cross-platform canine atlas and cross-species label transfer to human DRG. **(a)** Joint canine DRG atlas generated by integrating the deep full-length FLASH-seq dataset with an independently produced 10x Chromium single-nucleus DRG dataset. Colours indicate cluster identity according to the FLASH-seq annotations in this manuscript. Grey points represent cells from the independent 10x dataset. Names on the UMAP plot itself represent the consensus cluster names from the canine atlas. **(b)** Stacked bar plot showing results of cross-species label transfer from the NIH PRECISION human DRG consensus atlas, mapping FLASH-seq cell classes onto newly described human sensory neuron clusters.

In addition to the cross□platform canine integration, we further benchmarked our clusters against human sensory neuron diversity by performing Seurat label transfer from a recently generated multi□study human consensus atlas from the NIH PRECISION human pain network [10], which defines 22 neuronal classes. This analysis provided an independent, cross□species validation of our annotations and allowed us to identify the closest human counterparts for each cDRG subtype. Although some canine clusters—particularly within the C□nociceptor and proprioceptor groups—mapped onto multiple human subpopulations, the key neuronal types central to our ISH analyses (C□LTMR, PEP□KIT, SST/OSMR and non□peptidergic nociceptors) showed strong and unambiguous alignment with their human equivalents (Fig 5b). This convergence across datasets supports the conclusion that the cross□species comparisons underpinning our IL31RA and SSTR2 analyses rest on a conserved framework of sensory neuron identities.

## Discussion

Our study provides a deeply sequenced single-cell atlas of purified canine DRG neurons, generated through a FACS-based enrichment strategy combined with FLASH-seq. This dataset reveals the organisation of canine sensory neuron subtypes and establishes points of both similarity and divergence with human and mouse DRG.

### Sorting-based enrichment of neurons for FLASH-seq

Neuronal isolation from the DRG is challenging due to the predominance of non-neuronal cells. We report a neuron to non-neuronal cell ratio in canine tissue similar to that reported in human tissue[15], and significantly lower than that seen in rodents[3]. In human DRG, other investigators have used laser-capture based approaches to isolate individual neuronal somata to address this issue[86]. To address this here, we developed a FACS-based workflow that enriches for intact canine DRG neurons and preserves soma integrity. Although our current FACS parameters do not achieve clear size-based discrimination of large versus small neurons, our dissociation method maintains the full soma, unlike single-nucleus workflows. By measuring cell size with emerging imaging-enabled cell sorters[48], it may soon be possible to selectively enrich LTMR- versus nociceptor-sized neurons directly from flash-frozen post-mortem tissue. Such an approach could support targeted bulk-sequencing of DRG neurons from animals or human patients with defined painful conditions.

This study represents the first application of FLASH-seq to canine DRG neurons. Despite the modest number of sorted cells, the resulting dataset exhibits high quality transcriptomes, characterised by high unique gene detection and minimal mitochondrial content. Plate-based sequencing inherently captures fewer cells than droplet-based methods, and is therefore less effective for detection of rare cell clusters, but provides much greater transcriptional depth [32,43]. In our dataset, neurons generally exhibited a high number of detected genes, and this resulted in well-resolved clusters, even with fewer than 1000 cells sequenced. Nonetheless, some clusters were under-represented, particularly those corresponding to large-diameter neurons such as LTMRs and proprioceptors. Although we successfully detected these populations, they were likely biased against during FACS due to technical considerations.

We also observed a lower number of unique genes detected in tissue from female versus male dogs. We interpret these differences as potentially reflecting sex-dependent variation in membrane robustness or intracellular matrix properties ultimately affecting cell integrity during dissociation. Because sequencing depth varied between sexes, the present dataset cannot be used to draw meaningful biological conclusions about sex-dependent gene expression, despite this being an area of considerable interest in sensory neuroscience[29,31,55,71]. Nonetheless, clustering remained relatively stable between male and female-derived neurons, indicating that the overall transcriptional architecture is preserved despite these technical effects.

### A broadly conserved sensory neuron architecture across species

In our dataset, we identify cardinal classes of sensory neuron that are, perhaps unsurprisingly, similar to those described across mammals[11,42]. However, given the potential translational value of the dog, we were particularly interested in whether the canine DRG might be more like human rather than rodent ganglia. We first identified a proportion of non-neuronal cells in cDRG akin to that seen in humans. Given the growing literature surrounding the importance of neuroimmune interactions in pain[21,35,40,72], this comparative similarity in cellularity is a potential benefit when employing the dog for translation. We did not establish similarity at the level of specific immune cell populations, but others have recently found important commonalities between canine and human microglia-like cells, that are not mirrored in rodent DRG tissue[85].

Our results also point to an overall organisation of peptidergic nociceptors that is unlike that seen in rodents. Murine studies reveal clear separation among peptidergic nociceptor subtypes. Although terminology varies, these subtypes are commonly termed PEP 1.1/1.2/1.3/1.4, CGRP-Alpha/Beta/Gamma/Epsilon or CGRP + Sstr2/Dcn/Adra2a/Oprk1 [11,47,68]. In humans, these classes are less distinct and conflated into two classes[86]. Both our clustering and cross-species UMAPs show that canine peptidergic clusters are also less distinct according to the resolution afforded by our cell numbers. Beyond the peptidergic nociceptors, several canine marker genes resemble human rather than rodent patterns. CALCA, is more widely expressed across nociceptive populations in cDRG (see Fig S5), in particular in SST-expressing cells, as has also been shown in humans[69,70,86]. We also see limited canine expression of some classical marker genes, for example TH and SLC17A8 in C-LTMRs, with human marker genes such as CDH9 more highly expressed[10]. Furthermore, we identify a specific cluster of neurons, termed PEP0, identified previously only in human deep sequencing data[86]. PEP0 neurons have been suggested as a human specific population of A-fibre nociceptors, identified in both sequencing and Xenium spatial data[86], that express moderate levels of CALCA and PIEZO2 but do not have a clear function. In our data we see a similar population of neurons that meet QC thresholds but lack specific marker genes. Although these cells could simply represent low-quality neurons, we believe this to be unlikely due to the low percentage of mitochondrial reads and the observation that the cluster is still present when significantly higher QC thresholds are employed. To date, PEP0 type cells have only been identified in deep, plate-based sequencing datasets, and it is possible that their detection is biased against by more commonly used microfluidic sequencing methodologies.

Our data also integrates well with other sequencing datasets, further validating our clustering and highlighting more clearly the biological organisation of canine sensory populations. Combining our deeply sequenced dataset with another recent 10x-generated dataset[41], shows that canine cell types are generally analogous across technologies within species, and results in a collaborative resource with higher transcriptional coverage than each single study. Our data also aligns well across species to humans, with both single datasets[86] and consensus atlases[10] showing good agreement. This agreement is strongest in the case of the PEP-KIT neurons, non-peptidergic nociceptors, SST/OSMR neurons and C-LTMRs, that were the focus of our cross-species in situ experiments

### Rational cross-species therapeutic target identification

Our target-focused, cross-species comparison shows how gene expression differs in DRG neuron subtypes across species and hence how this may affect translation of therapeutics. This analysis identified a set of highly conserved genes across dog, human and mouse DRG, including OPRM1, NTRK1, and SCN10A. These targets show conserved, subtype-specific enrichment across all three species, supporting the view that these core nociceptor signalling molecules are likely to be pharmacologically tractable across dogs, humans and rodents. Consistent with this, each of these three genes is already the basis of successful analgesic therapies in clinical or veterinary clinical use[27,65,80].

However, divergence in the expression of other potentially therapeutically relevant genes highlights important species-specific specialisations. We focused on two examples, IL31RA and SSTR2, where cross-species differences may have direct implications for both drug mechanisms and genetic access strategies.

In the case of IL31RA, we observed a strikingly restricted neuronal expression pattern in the dog, where IL31RA expression is confined to the SST/OSMR population. In contrast, human DRG shows a distributed pattern where IL31RA is expressed across nonpeptidergic nociceptors and CLTMRs in addition to SST neurons. This was confirmed through *in situ* hybridisation in both species. These neuronal subtypes possess distinct spinal projections and engage different downstream circuits[24,56,62,78]. Therefore, this divergence suggests that IL31–targeted therapies may modulate a qualitatively different itch percept in dogs and humans. Extending beyond circuitry, IL31RA has previously been identified as a domestication associated gene[60], and its reduced abundance in dogs raises the possibility, albeit speculative, that selection pressures related to a desire to increase parasite tolerance during cohabitation may have shaped IL31–dependent itch pathways. This theory is consistent with the finding that dogs do not scratch consistently in response to many common experimental pruritogens[6,19]. Interestingly, IL31RA agonism has recently been reported to act as an immunological brake, reducing the severity of skin inflammation in pruritic conditions[28]. It is therefore possible that comparatively reduced IL31RA expression could also be a factor underlying high prevalence of atopic dermatitis in dogs[26].

In the dog, we find SSTR2 to be expressed across a broad set of sensory neuron subtypes, spanning small-diameter nociceptors through to medium diameter neurons and even some LTMR-like cells. This contrasts with the situation in rodents, where SSTR2 marks a discrete nociceptor subclass recently implicated in spontaneous neuropathic pain[88]; making those cells appealing for targeted therapies. Importantly, this divergence likely reflects changes in marker gene identity, not the absence of the underlying cell class, but nonetheless has substantial implications for gene targeting approaches that require molecularly specific access points[47,53]. Whether the human SSTR2 pattern more closely resembles dog or mouse remains unresolved, although it appears that SSTR4 is a more appealing analgesic target in humans with the potential for fewer endocrine side effects[14,75]. Interestingly, SSTR4 has not yet been identified in the canine genome, and this underscores the need for continued comparative profiling[39]. Together, these examples illustrate that while many key sensory targets are conserved, species specific transcriptional features may affect both the mechanisms and the translational relevance of pain and itch therapies.

In conclusion, our results establish a high-resolution resource and demonstrate that deep, full-length profiling of purified cells from frozen post-mortem tissue is feasible. Using these data, we outline the conserved and divergent, and in places potentially domestication shaped, sensory pathways that refine the dog’s translational value for pain and itch.

## ACKNOWLEDGEMENTS

The work was supported by a PhD studentship grant from the University of Glasgow Vet Fund. A.M.B is supported by a Wellcome Trust Early-Career Award (304005/Z/23/Z), and G.A.W is supported by an MRC grant UKRI3822.

We are grateful to the Scottish SPCA and Charles River Laboratories for their kind assistance with sourcing canine tissues for this project. We acknowledge the expert assistance of Daniel Lewis and Diane Vaughan, MVLS-SRF Flow Cytometry Core Lab at the University of Glasgow. We are grateful to Andrew McCluskey and Joy Kabagenyi for their guidance in analysis of the single-cell dataset and to Matthew Sapio and Michael Iadarola for their assistance with reanalysis of their existing bulk sequencing data. We also thank Leanne Jankelunas, Shamsuddin Bhuiyan, William Renthal, and Rell Parker for their collaborative role in generating the canine 10x atlas and for their continued engagement, discussion and assistance during the preparation and submission of this and the companion manuscript.

Raw sequencing data and processed files including the Seurat object are available at the Gene Expression Omnibus (GSE327952). An interactive browser of FLASH-seq data is available at https://canineflashseq.shinyapps.io/Browser/. Upon publication of the manuscript, all other data and analysis files will be freely available at the University of Glasgow’s Enlighten repository.

The authors declare no competing financial interest.

For the purpose of open access, the author has applied a Creative Commons Attribution (CC-BY) licence to any Author Accepted Manuscript version arising from this submission.

