## Supplemental material for "Deep FLASH-seq profiling of purified canine sensory neurons uncovers species-specific signatures relevant to pain and itch"

Contains Supplemental figures 1-10 & Tables S1, S2, S3, S5 & S6

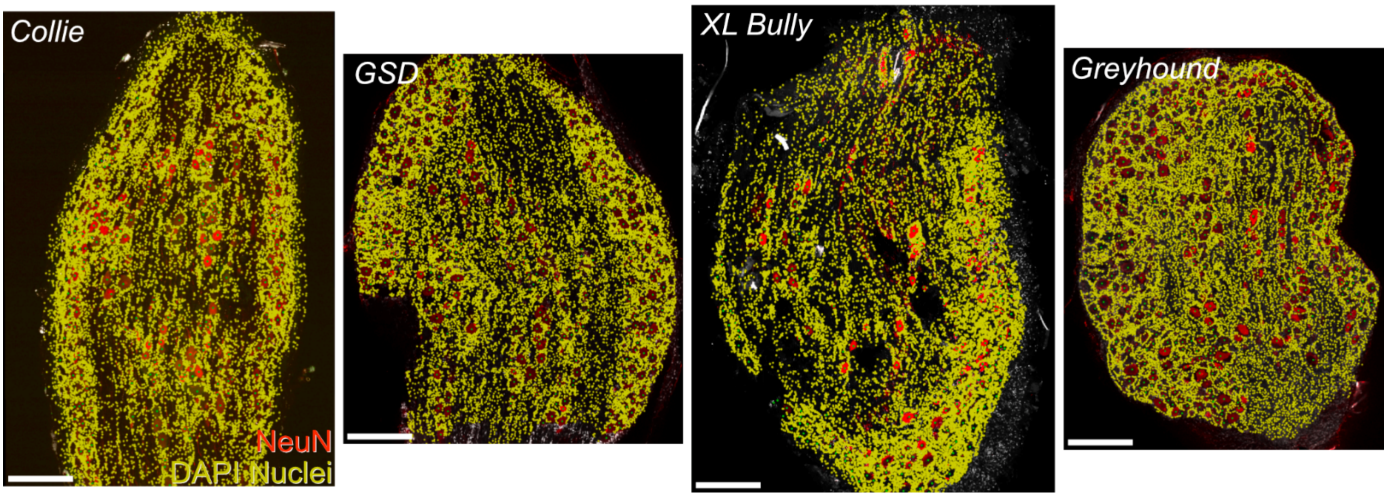


**Fig S1**. Example confocal images of cDRG from 4 animals in which the ratio of neurons to non-neuronal cells has been estimated. Neurons are stained with NeuN and all nuclei stained with DAPI. Each yellow dot annotation represents a single counted non-neuronal nucleus, while each green dot represents a counted neuron. Images are single confocal sections acquired at one airy unit with a 20x objective. Scale bars represent 400µm.

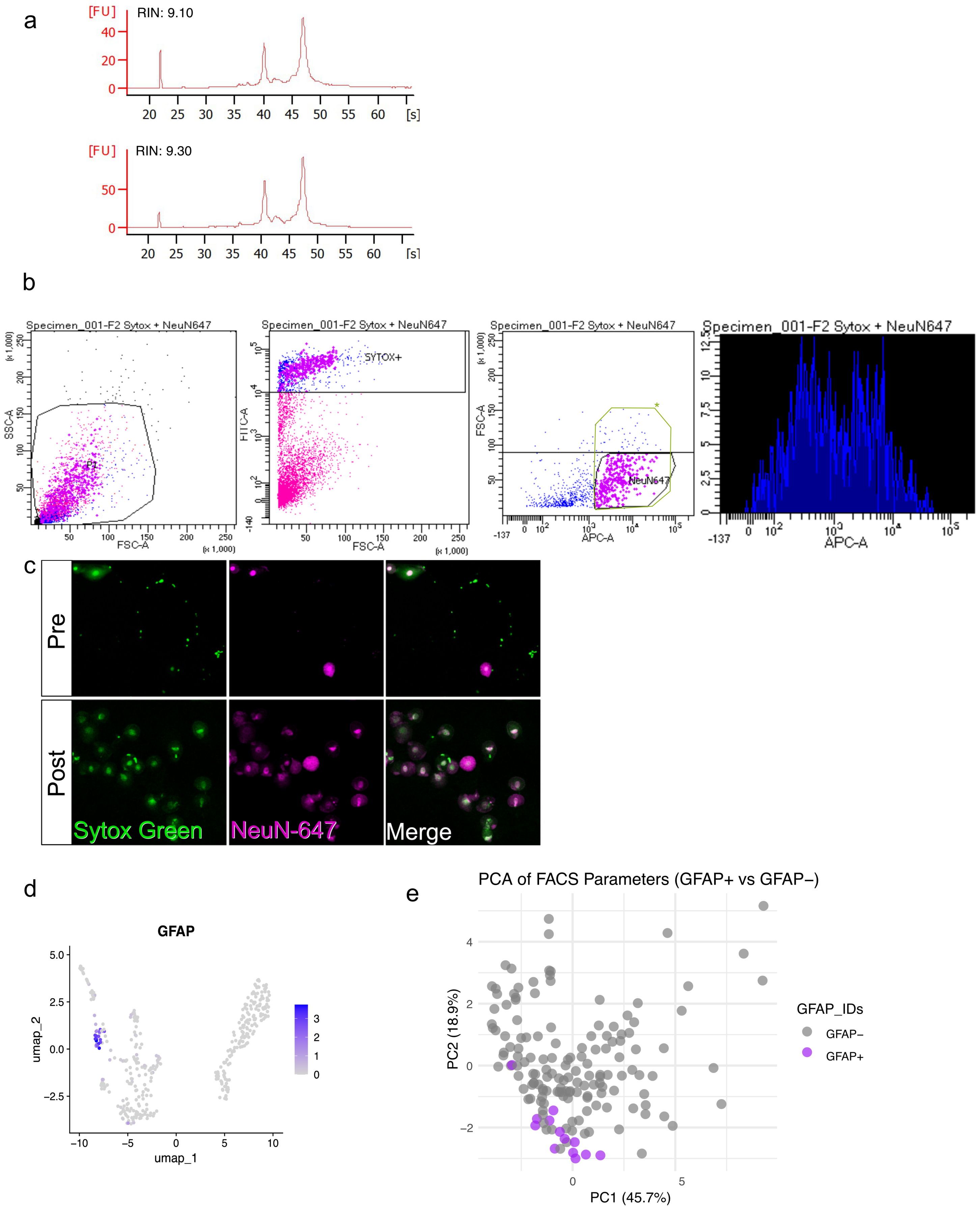


**Fig S2.** **(a)** Example RNA Integrity Scores (RIN) of DRG collected from dogs submitted for sequencing. **(b)** FACS gates shown in order (left to right) to achieve neuronal isolation for plate-sorting. SSC-A vs FSC-A gate scatter gate differentiated small debris from putative cells; FITC-A vs FSC-A gate was used to identify cells with high SytoxGreen fluorescence; FSC-A Vs APC-A gate identified NeuN647 positive cells (neurons). Refinement of Neun647 gating was aided by 1-parameter histogram of the APC-A fluorescent channels, X axis is APC-A and Y axis is number of events (counts) for each fluorescent level. Events included after the inflection point (observed at $10^3$ in this graph) were sorted into a plate. FSC-A threshold was informed by which FSC-A level filtered out complex cells formed by neurons with immune cells attached on its surface. To do this we isolated cells at different FSC-A levels followed by observation under the microscope. Notice that FSC-A upper threshold was reduced from 150000 ( green square, *) to 100000 between trial and definitive plates due to high of number of well with no sequencing outcome resulting from events above this threshold. **(c) Top -Pre FACS cell suspension.** Representative fluorescence image after DRG dissociation but prior to FACS. Nuclei were labelled with SytoxGreen to visualise all nucleated events; NeuN647 staining highlights neuronal cells. Notice the heterogeneous mixture of neuronal cells and non-neuronal nuclei. **Bottom-Post FACS cell suspension**. Fluorescence image of the sorted population gated for SytoxGreen+/Neun647+ cells. Post-sort fractions show a visible enrichment for neuronal cells and preservation of neuronal cell integrity. **(d) Glial cells (GFAP+) cluster seen in Trial Plate FeaturePlot.** Presence of a distinguishable GFAP cluster evidenced minor glial cell contamination during FACS sorting. GFAP expression in glial cells, allows removal of this cluster for downstream analysis. **(e) PC1 Vs PC2 scatter plot from principal Component analysis (PCA) reveals that it is not possible to differentiate GFAP cells from neuronal cells through FACS parameters alone.** PCA analysis using all FACS parameters was performed to assess if GFAP cells were distinguishable from non-GFAP cells during FACS. Each purple dot in the scatter plot represents a GFAP-positive cell, and each grey dot corresponds to a non-GFAP cell. The first two principal components accounted for a substantial proportion of the total variance, PC1 explaining 45.7% and PC2 explaining 18.9% of the observed variability. However, the overlapping distributions indicates that the measured features included in the PCA are insufficient to reliably distinguish these two cell populations.

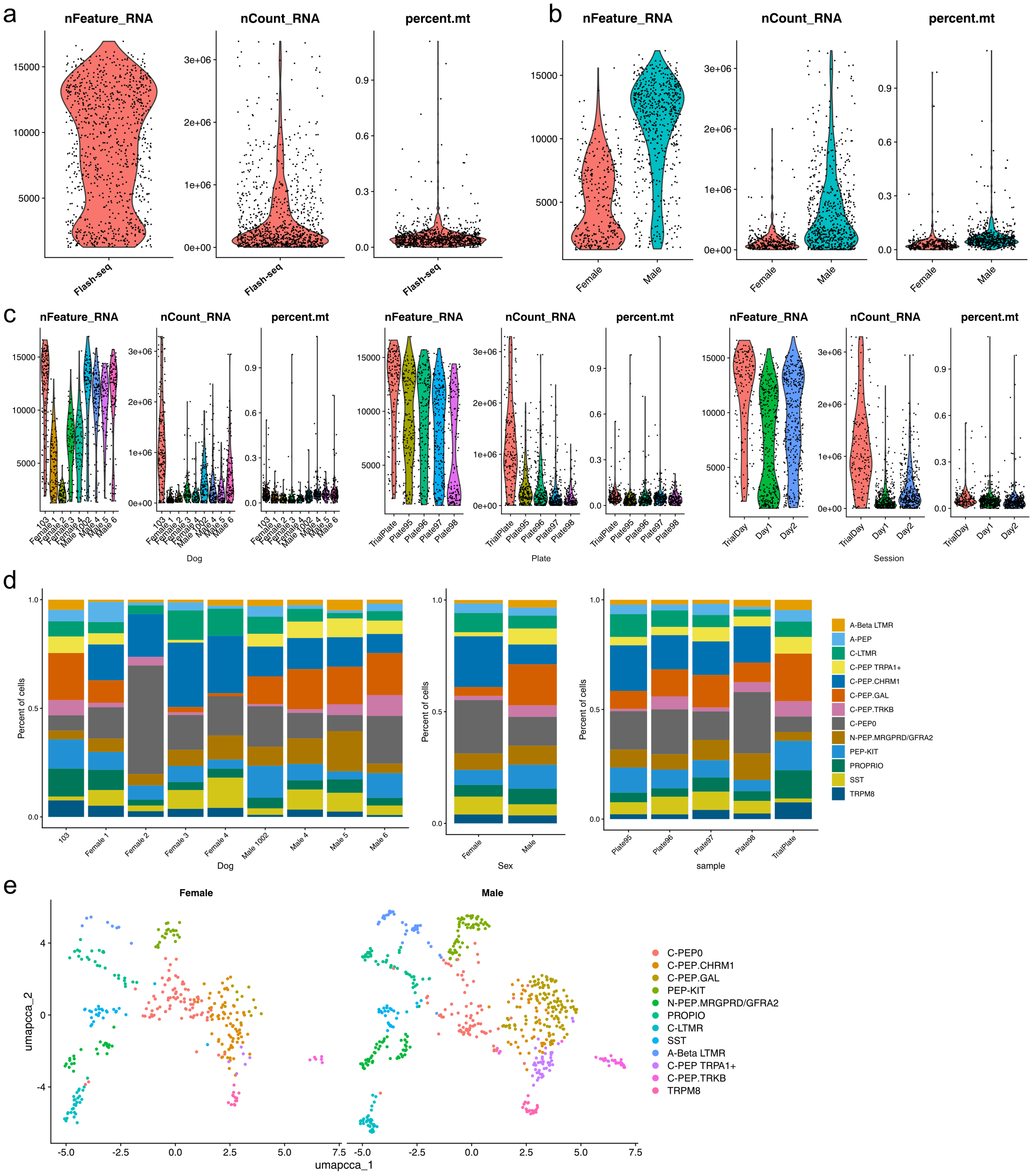


**Fig S3. Quality control metrics for FLASH-seq sequencing data. (a)** Violin plots showing number of unique genes per cell (nFeature_RNA), total per cell counts (nCount_RNA) and mitochondrial gene percentages (percent.mt). **(b&c)** Violin plots showing these same QC metrics across sexes, animals, plates and sorting sessions. **(d)** Stacked bar plots of final cluster proportions across animals, sexes and plates after CCA integration for batch correction. **(e)** UMAP plots showing final clusters separated by those seen in male and female animals after CCA integration.

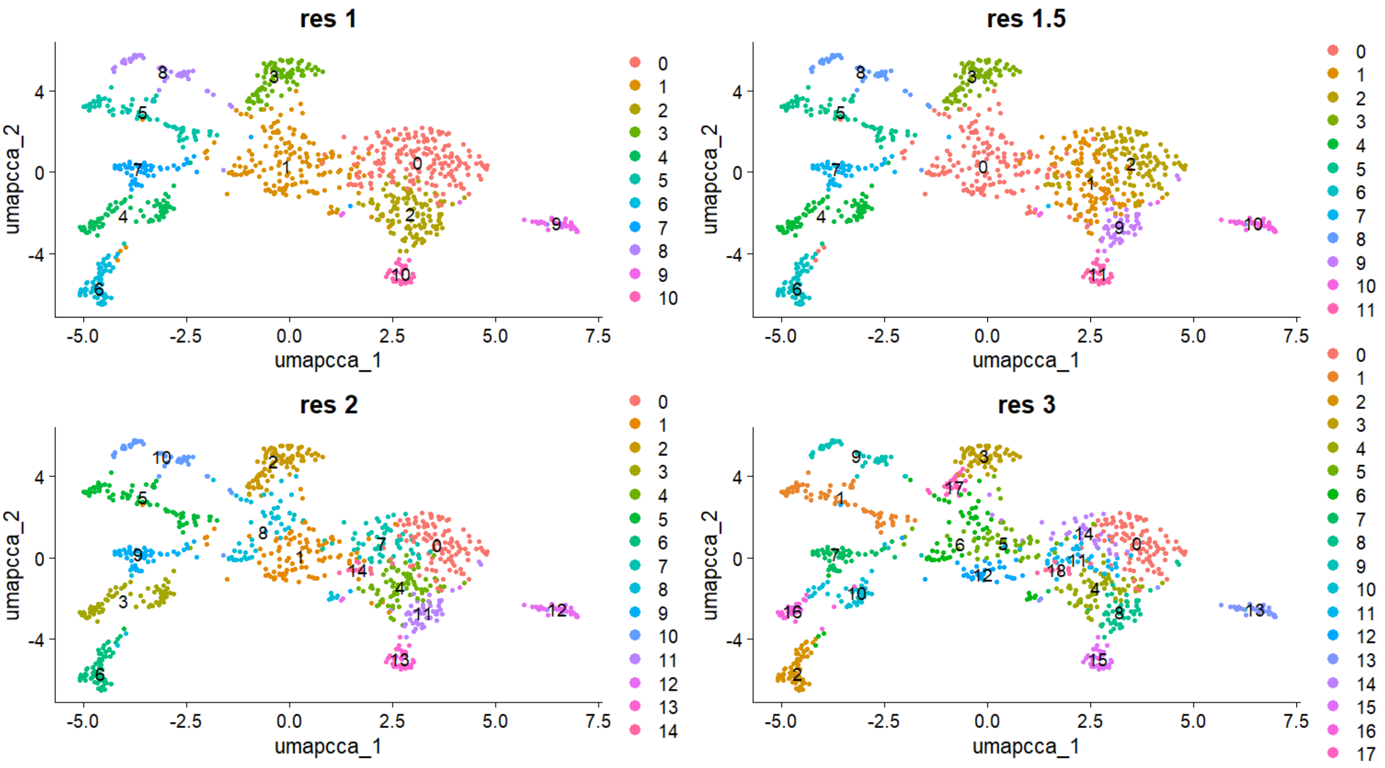


**Fig S4. Results of different cluster resolutions on Seurat clustering.** A resolution of 1.5 was used for the final dataset.

a. DRG Neuronal subtype marker genes from Jung et al. 2023

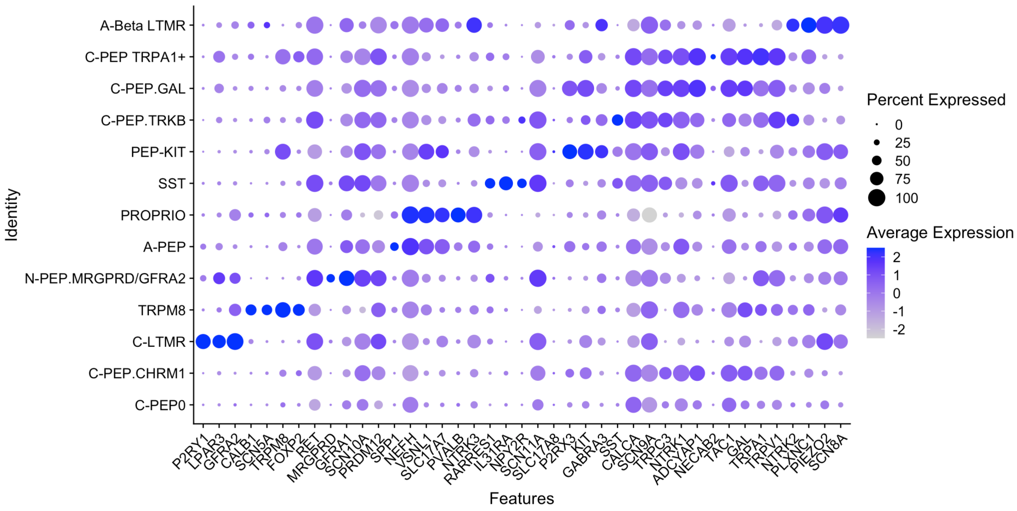


b. DRG Neuronal subtype marker genes from Bhuiyan et al. 2024

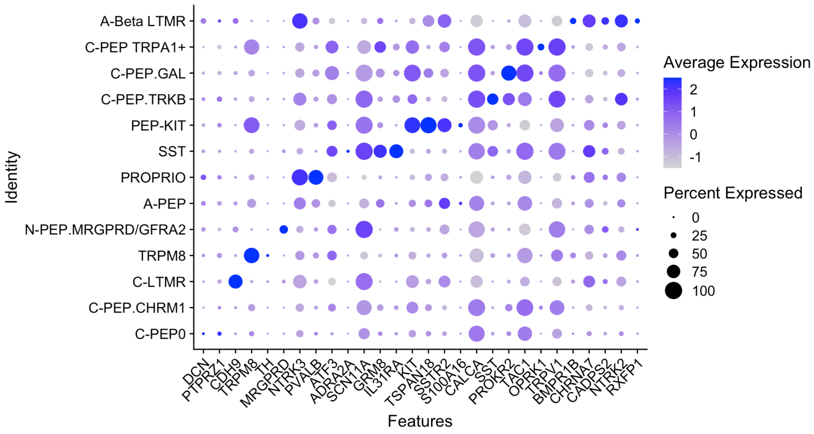


**Fig S5. Marker genes from other cross-species atlases**. Expression shown across canine FLASH-seq clusters. Dot colour represents scaled mean expression; dot size represents the percentage of expressing cells.

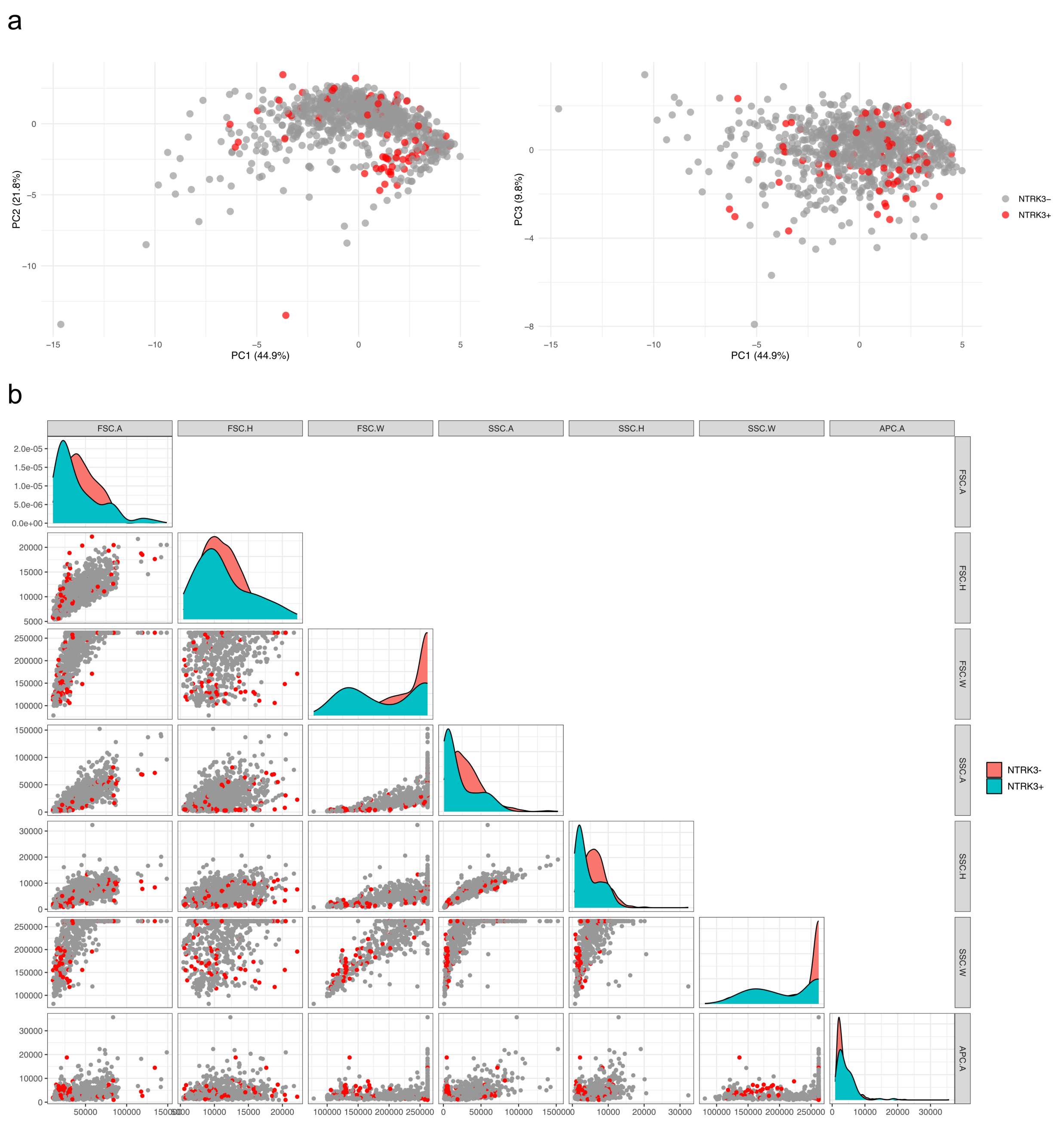


**Fig S6. Non-distinguishable nature of large vs small diameter neurons using FACS parameters. (a)** Principal component analysis of FACS parameters (scatter and fluorescence parameters) with large (NTRK3+) and other cells indicated by colour. Putative large cells overlap significantly in the principal component space, indicating that FACS parameters alone cannot be used to separate these cells. **(b)** Pairwise scatter plots of FACS scatter gating parameters, again with NTRK3+ cells indicated in red. Although large cells appear to show reduced scatter, they are not separable from other cells.

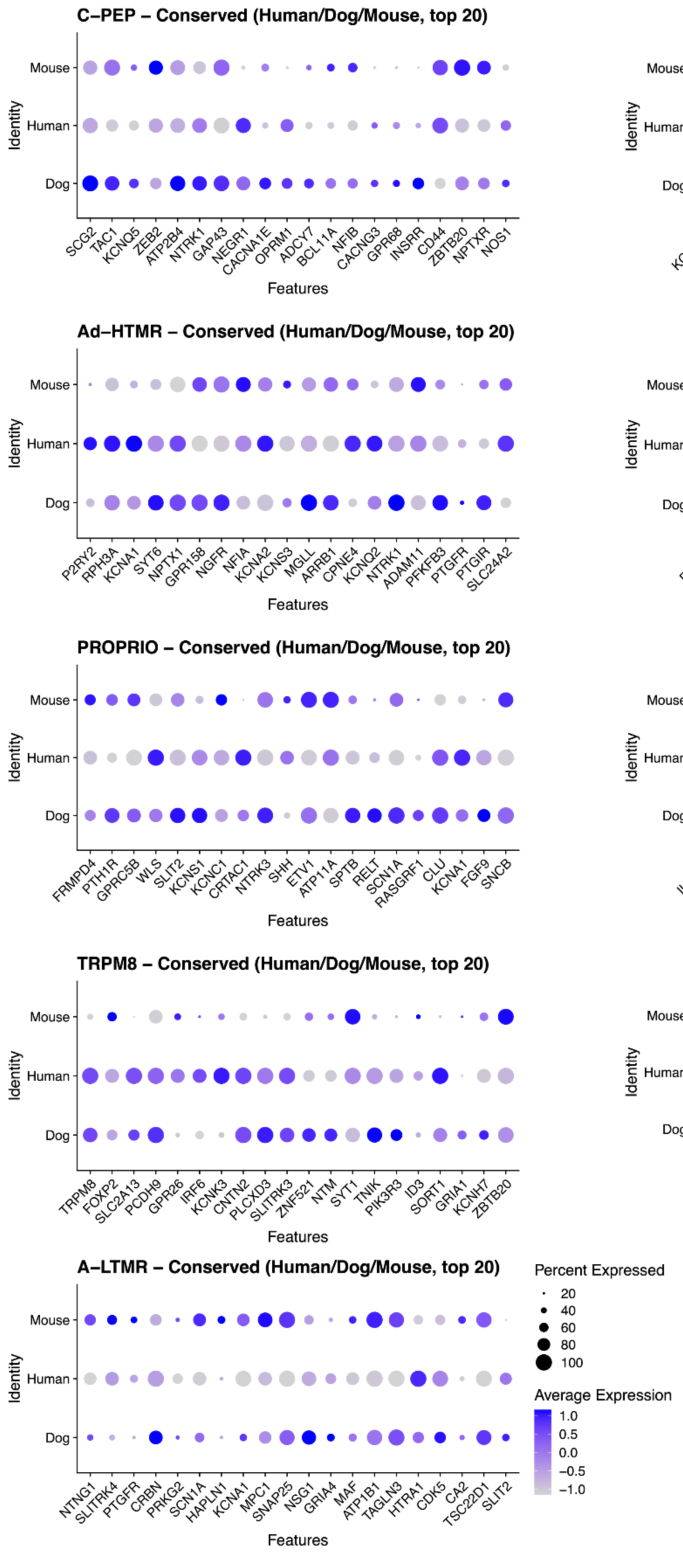


**Fig S7. Cross-species convergent expression patterns of functional genes across unified neuronal clusters.** Dot plots showing expression of all genes belonging to major functional categories (transcription factors, GPCRs, ligands, ion channels, synaptic proteins) across the unified dog, human, and mouse DRG clusters. Dot colour represents scaled mean expression; dot size represents the percentage of expressing cells. For each broad class of sensory neuron, the top 20 convergent genes are shown. These were identified using FindConservedMarkers, retaining only genes significantly enriched in that cluster in all three species ($\log2FC \geq 0.25$, ≥10% expressing, adjusted $P < 0.05$) and ranking them by the maximum adjusted P value across species.

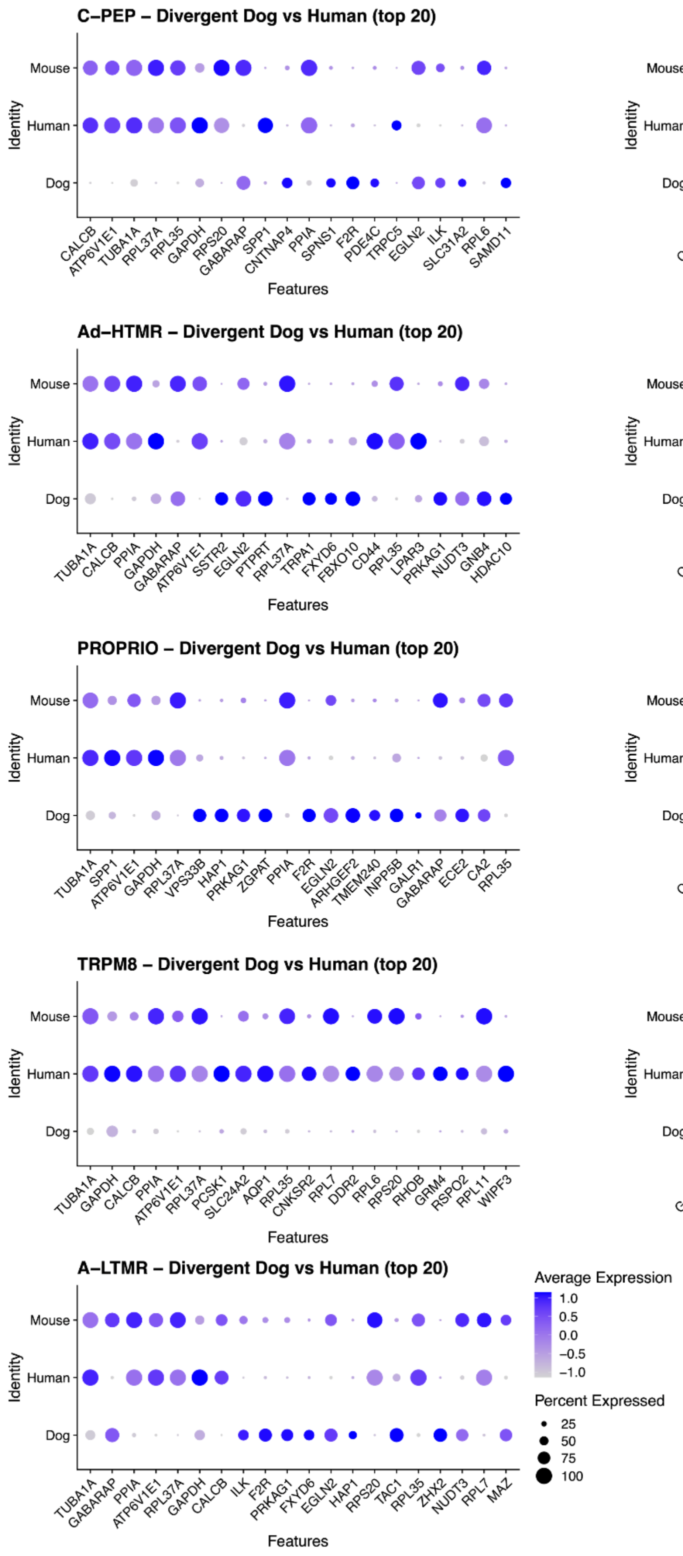


**Fig S8. Divergent gene expression between dog and human across unified neuronal clusters.** Dot plots showing the top 20 genes with significant species-biased expression between dog and human within each unified cluster; dot colour represents scaled mean expression and dot size the percentage of expressing cells. The top 20 divergent genes were selected by identifying genes significantly enriched in either dog or human (log2FC ≥ 0.25, ≥10% expressing, adjusted P < 0.05) and ranking them by adjusted P value within the species-bias test.

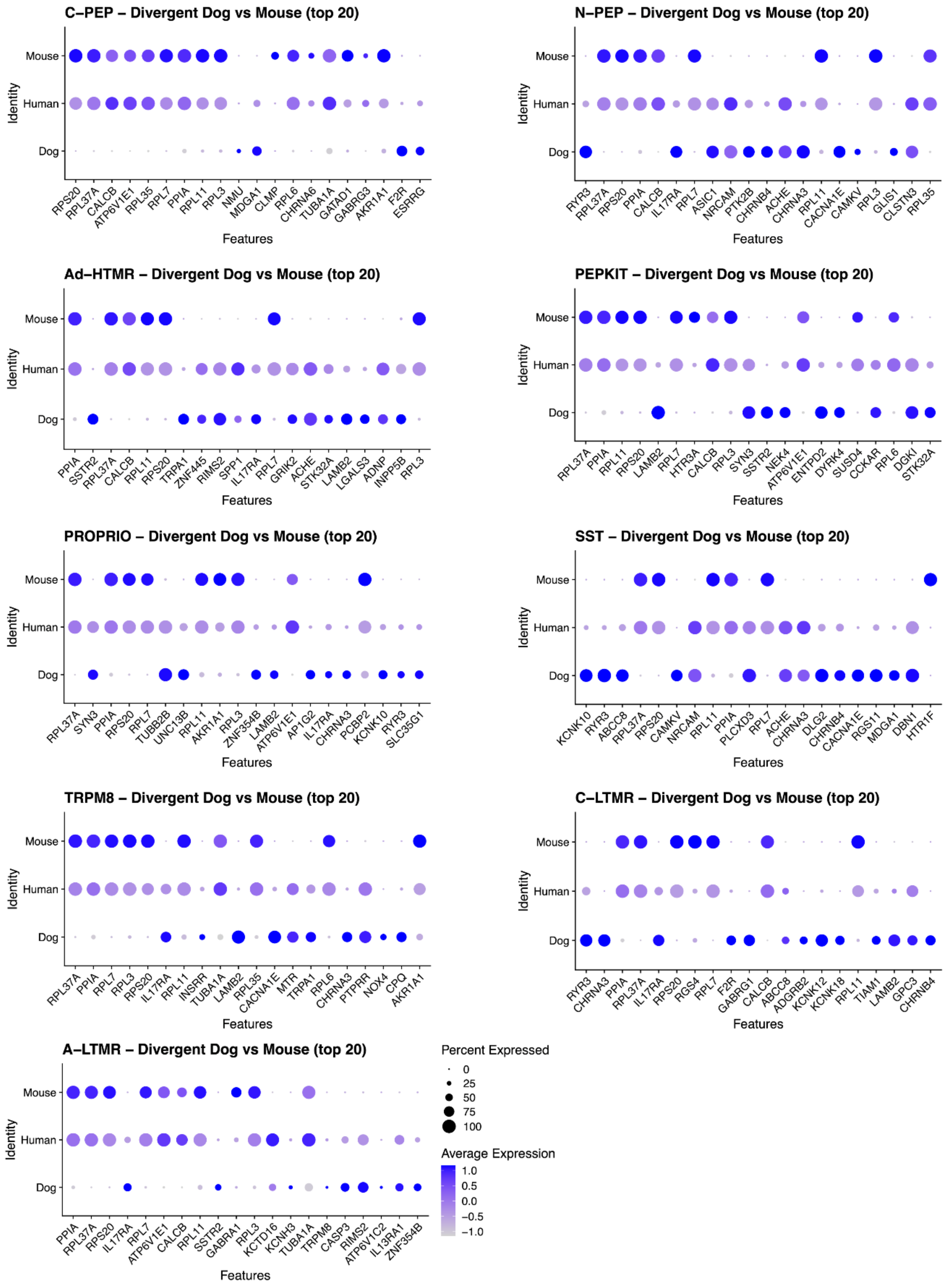


**Fig S9. Divergent gene expression between dog and mouse across unified neuronal clusters.** Dot plots showing the top 20 genes with significant species-biased expression between dog and mouse within each unified cluster; dot colour represents scaled mean expression and dot size the percentage of expressing cells. The top 20 divergent genes were selected by identifying genes significantly enriched in either dog or mouse (log2FC ≥ 0.25, ≥10% expressing, adjusted P < 0.05) and ranking them by adjusted P value within the species-bias test.

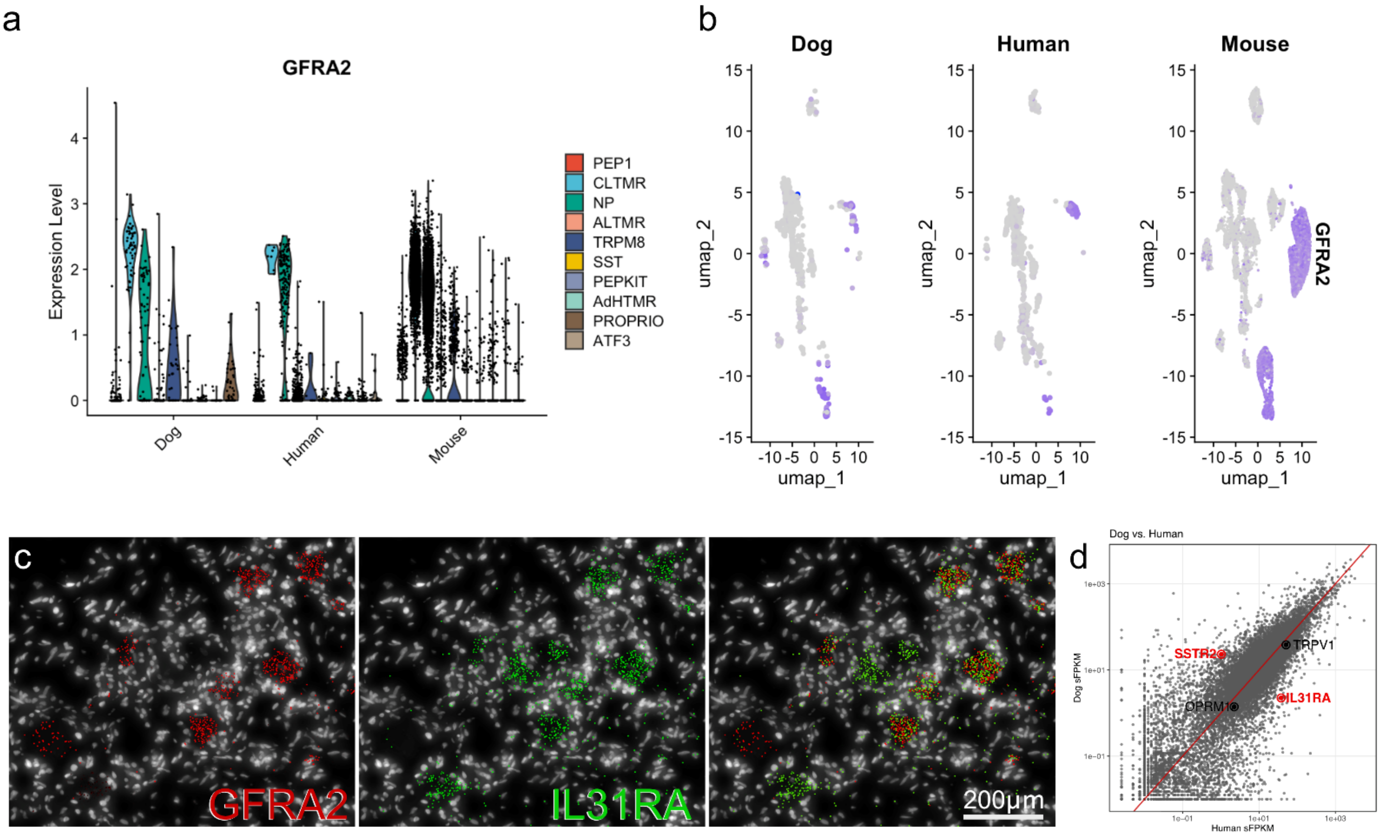


**Fig S10. Data to support in situ analyses. (a)** Violin plots showing *GFRA2* marker gene expression across unified clusters in dog, human and mouse. **(b)** UMAP feature plots showing the same data and confirming that high *GFRA2* expression occurs in non-peptidergic and C-LTMR clusters and not in SST/OSMR neurons across species. **(c)** Image from Xenium in situ platform using human DRG data from Yu et al. (2024). *GFRA2* and *IL31RA* transcripts are shown in red and green, respectively, with DAPI nuclei in white. Reanalysis of this data demonstrates a significant overlap in cellular expression between the two genes in human DRG tissue. **(d)** Reanalysis of bulk sequencing data from Iadarola et al. (2018) where transcript abundance in human DRG bulk-seq data is compared to that in a high-quality canine dataset. An excellent correlation is evident, but some outliers are seen. On this basis *IL31RA* transcript abundance is lower in canine DRG than in humans whereas *SSTR2* is higher.

| mRNA | Gene name | Organism | Nº of binding sites | Probe Channel | Amplifier |
|---|---|---|---|---|---|
| IL31RA | Interleukin 31 Receptor A | Dog (*Canis lupus familiaris*) | Up to 20 | Channel 3 | Alexa Fluor-647 |
| GFRA2 | GDNF family receptor alpha 2 | Dog (*Canis lupus familiaris*) | Up to 20 | Channel 1 | Alexa Fluor-546 |
| SSTR2 | Somatostatin receptor 2 | Dog (*Canis lupus familiaris*) | Up to 20 | Channel 2 | Alexa Fluor-546 |
| KIT | KIT proto-oncogene receptor tyrosine kinase | Dog (*Canis lupus familiaris*) | Up to 20 | Channel 1 | Alexa Fluor-647 |

**Table S1. Summary table of HCR™ Gold RNA-FISH probes and amplifiers.**

| Breed | n | Mean | Median | sd | Min | Max | Q1 | Q3 |
|---|---|---|---|---|---|---|---|---|
| Collie | 100 | 47.8 | 46.3 | 15.0 | 23.5 | 80.9 | 35.8 | 58.7 |
| GSD | 100 | 48.7 | 49.0 | 15.2 | 22.3 | 81.0 | 35.7 | 59.6 |
| Greyhound | 100 | 49.1 | 49.2 | 14.1 | 21.1 | 81.0 | 38.2 | 60.5 |
| XL Bullie | 100 | 44.7 | 42.0 | 13.7 | 21.0 | 80.4 | 33.6 | 54.8 |

**Table S2. Individual dog data used for neuronal size analysis in Figure 1.** Data represents diameters in μm.

| Dog ID | Dog Breed | Age (year, y; months, m) | Sex | RIN (if applicable) | Technique |
|---|---|---|---|---|---|
| Greyhound | Lurcher X | 2y | Male | | IHC |
| German shepherd | German shepherd | 2y | Male | | IHC |
| Collie | Collie X | 2y | Male | | IHC |
| Bulldog | Bulldog X | 2y | Male | | IHC |
| 1001 | Beagle | 1y | Male | 9.3 | FLASH-seq |
| 1003 | Beagle | 1y | Male | 10 | FLASH-seq |
| Male 1002 | Beagle | 1y | Male | 9.1 | FLASH-seq |
| Male 4 | Beagle | 2y | Male | 9.3 | FLASH-seq |
| Male 5 | Beagle | 2y | Male | 9 | FLASH-seq |
| Male 6 | Beagle | 2y | Male | 9.7 | FLASH-seq, FISH |
| Female 1 | Beagle | 9m | Female | 8.8 | FLASH-seq |
| Female 2 | Beagle | 9m | Female | 7.8 | FLASH-seq |
| Female 3 | Beagle | 1y | Female | 8.4 | FLASH-seq |
| Female 4 | Beagle | 1y | Female | 8.7 | FLASH-seq |
| Male1003 | Beagle | 9m | Male | | FISH |
| FC1 | Beagle | 1y | Female | | FISH |
| F2 | Beagle | 10m | Female | | FISH |

**Table S3. Individual information for each dog:** Dog ID, breed, sex, age, use (immunohistochemistry, IHC; in situ hybridization, ISH) and RNA Integrity Score (RIN).

**For Table S4** see additional spreadsheet

| | Number Small (<40µm) Neurons | Percentage of Small (<40µm) Neurons | Number Medium (40-60µm) Neurons | Percentage of Medium (40-60µm) Neurons | Number Large (>60µm) Neurons | Percentage of Large(>60µm) Neurons |
|---|---|---|---|---|---|---|
| **SSTR2** | | | | | | |
| SSTR2- | 76.25 (65 - 107) | 57% (49% - 67%) | 59 (40 - 76) | 54% (44% - 69%) | 20.25 (9 - 32) | 72% (53% - 79%) |
| SSTR2+ | 38.25 (33 - 46) | 29% (27% - 30%) | 26 (19 - 30) | 24% (20% - 33%) | 4.25 (2 - 7) | 15% (8% - 29%) |
| SSTR2++ | 18.75 (6 - 30) | 14% (4% - 22%) | 23.5 (8 - 41) | 22% (7% - 28%) | 3.5 (2 - 5) | 13% (9% - 18%) |
| **KIT** | | | | | | |
| KIT- | 51.75 (33 - 72) | 39% (28% - 45%) | 88 (69 - 112) | 81% (77% - 92%) | 27.75 (14 - 42) | 99% (98% - 100%) |
| KIT+ | 38.75 (30 - 48) | 29% (25% - 41%) | 12.5 (3 - 20) | 12% (3% - 16%) | 0.25 (0 - 1) | 1% (0% - 2%) |
| KIT++ | 42.75 (35 - 52) | 32% (28% - 38%) | 8 (5 - 14) | 7% (6% - 10%) | 0 | 0% |

**Table S5.** Neuronal counts (shown as mean and range) for SSTR2 and KIT categories per neuronal size class. Data is from 4 animals. Neuronal diameter categories grouped in small (<40 µm), medium (40–60 µm) and large (>60 µm). Data used for Figure 4.

**Table S6.** Full list of genes from domestication studies overlapping with cluster DEGs

| Cluster | Gene |
|---|---|
| A-Beta LTMR | NTNG1, ZACN, DEFB119, ZMAT4, TMEM26, GALR2, SSH2, BTAF1, CEP135, TMC8, UNC13B, CACNA1A, BEND5, C5AR1, EXOC7, FHOD3, TIAM1, FSTL4, KIAA0586, FANCG, ZNF516, PDE7B, SEMA4A, POLR1E, DNAH17, ARHGEF39, KIAA1328, COPS7B, SCN2A, ACAD11, REEP3, ACOX1, DNMBP, IDE, PIGO, MAPKAPK3, LMBR1L, LYST, DNAH3, PLOD2, ACSM5, ZNF236, TTC39A, RYR3, DUT, NRG2, ELF2, CYFIP1, RALGAPA2, SBF2, SUGP1, PCSK4, MARK3, CADM2, TNKS2, BACH2, ADM, CAMKMT, ITCH, HECW2, ZZEF1, TMEM120B, PIK3R1, ATXN2, POMT1, ART3, MDM2, FKBP11, XYLB, SF3B1, LARS2, CNTN5, MAU2, NUP54, ARHGAP32, LRRTM3, BAZ2B, SI, VEZT, DIP2C, LARP1B, LRIG3, RNPC3, AHCYL2, NUP107, USP15, TUBGCP5, NR2C1, CNOT2, HYDIN, TRMT1, SCYL3, MTMR4, AFMID, STX10, DNAL1, ZNF396, CELF1, RFX3, AQP11, NPHP3, ROR1, PLXDC2, MYCBP2, PLEKHH2, STK10, CALB2, TOM1L2, FGD6, LMOD1, RFXANK, CMTR2, RHOF, MON2, ZNF24, ACVR1, ZNF407, PIGL, PIK3C3, GBF1, ZNF23, POLN, HTR2B, MAP6, HERC2, MFSD8, GALR1, CCNT1, PRKN, ST7, CPEB3, ASIP, PMM2, ARID1B, CNDP1, EHHADH, PAQR6, TRDN, CEP78, MBD2, SEPTIN4, RANBP17, PPM1H, PSD2, POLI, COL11A1, ADCY8, WDR26, RMDN2, KXD1, IKZF1, MET, ANGPTL7, SMCHD1, NRXN3, KLF12, RAB3GAP1, ADCY6, FRG1, NAT8L |
| A-PEP | ZMAT4, NAT8L, NGFR, HAPLN4, SEMA3D, ADM, ADCY8, RNF6, GALR1 |
| C-LTMR | RTBDN, DAPL1, FRMD6, TLX3, SMO, FGF1, CACNB3, ABHD12, BACH2, CRYM, GALR1, SAMD12, NEDD4L, C3orf18, TSPAN14, GDPD5, KCNC2, MAPKAPK3, LIMCH1, METTL22, LMNA, REEP6, CUX2, RALY, GRB10, CISH, NAV1, ZC3H12A, CNTN5, ADAMTSL5, DOCK2, TBC1D22A, PSAT1, GAD2, METAP2, TIAM1, PCSK4, ASAP1, NTAN1, CTH, BDH2, PLCXD3, PRMT2, PLEKHA4, MEIS3, RFXANK, ACVR2B, GLIS1, CEP78, ACOX1 |
| C-PEP TRPA1+ | MYOF, FBN1, NCAM2, GPR139, SCUBE1, PLCXD3, FOXP2, NEDD4L, KCNJ3, PRRC2B, FAM107B, CYP1B1, PCDHGC5, PRKACB, FKBP8, PYGB, PNMA1, RIMS2, PRMT2, TLN1, CARHSP1, YWHAH, DIP2C, CACNB3, ACOT6, EEF1A1, HSPD1, ELAVL4, CNDP2, EPS15, CCSER2, SLCO2B1, SUSD6, PSAT1, COL11A1, GNAQ, RFTN2, GDPD5, PEMT, TSPAN14, MOB4, MTOR, HSD17B12, RPS15, PRDX3, CCT3, GNA12, SNX19, ZFYVE28, RNF103, SERPINH1, NKAIN2, ANKRD44, TMEM59L, SNRPN, SETD9, PCDH18, PLEKHM3, TOMM20L, ZMYND11, NPM1, OPRM1, SRP72, PGRMC2, ATG13, ASAH1, GRB10, CETN3, ACTR10, ABHD12, LRPPRC, PSMD1, ACSS2, FBXL3, TLX3, RRN3, SLC25A44, ARRB1, VCP, GTF2E2, ARL2BP, PARP12, SDAD1, UCK1, NINL, CUEDC1, NIPA2, RPS3, PRNP, COQ10B, CALR, CUX2, BCAT2, HARBI1, GNL2, PPM1H, HPD, NCAPG, ARID4A, PSD4, REEP1, SMIM15, FARSA, AMBRA1, CREB3, NAT8L, DDX23, MRPS23, MARS2, DHX9, KAT7, PDXDC1, LMNA, MANEAL, METAP2, NTAN1, ITGB1, HIPK2, RNF6, PGS1, TVP23B, ARID1B, SNIP1, PPM1E, RAD23A, FRMPD1, SMOX, DIAPH1, SMCHD1, PIK3C3, POLI, STARD6, HINT2, CNIH4, CLN5, TBC1D22A |
| C-PEP.CHRM1 | FBN1, NCOR2, TOPORS, FRG1, BAZ2B, DNASE2, SRSF11, RYR3, NRXN3, COL11A1, SCN2A, UNC13B, ACVR2B, ELF2, RNPC3, ARID4A, CDC42BPA, MED13, MYCBP2, NCOA6, GPR139, HERC2, GDPD5, IKZF1 |
| C-PEP.GAL | CLIC4, FKBP8, FLVCR2, EEF1A1, HINT2, SCUBE1, GTF2E2, MOB4, PRDX3, RPS15, UCK1, PNMA1, YWHAH, TLN1, CNN3, OPRM1, GNAQ, SYN2, PCDH18, ACTR10, PSMD1, PCDHGC5, RIMS2, NCAM2, EPS15, CARHSP1, PRRC2B, HSD17B12, NPM1, HSPD1, PLEKHM3, PRMT2, NGFR, LCORL, RPS3, COQ10B, DNAJC15, ZMYND11, ABAT, LRPPRC, UBA52, PLCXD3, CCT3, ARRB1, NIPA2, FBXL3, CCDC107, GADD45GIP1, ELAVL4, CUL1, ASAH1, MANEAL, RNF103, PGRMC2, PTPMT1, ACOT6, CNDP2, REEP1, NXPH3, CAPS2, PYGB, KAT7, GNA12, RBPMS, CETN3, SETD9, RAP1B, SNX19, PEMT, PDE6D, PACRG, VCP, NCAPG, GCDH, RAD23A, SYNGR2, CREB3, ARL2BP, RNF6, LMNA, EHHADH, FXR1, PRKACB, SNRPN, STARD6, SLC35B1, KCNJ3, PAK1, SRP68, SPOP, TMEM59L, GLMP, EIF2S2, PPM1E, FRMPD1, CNIH4, SLC44A3, MXD4, CALR, SLC25A44, TLX3, CISH, SRP72, RAI1, BCAT2, ATG13, DNAJC19, FARSA, TRMT61A, ZFYVE28, UBA5, NUBPL, AGAP1, TBC1D22A, NKAIN2, ARID4A, DDX23, MRPS23, TVP23B, PSD4, SLK, GBA2, RFTN2, PSAT1, COMMD10, HPD, CUEDC1, RSPRY1, EXOC7, SMIM15, PDXDC1, COG6, METAP2, UBIAD1, RANBP17, PGS1, CCSER2, MAP6, LRRN3, KBTBD4, CACNB3, RAD51C, ANKS4B, NOCT, RCN2, GPR139, TMEM167A, ITGB1, CELF1, ABHD12, DCAF10, ACSS2, NACC1, CNOT4, FAM169A, EXOSC3, MEAF6, DHX9, IPO9, FAM117A, GNL2, SUSD6, TPGS2, NR2C2AP, JRKL, ARMC9, AMBRA1, RND1, FBXW11, NVL, HARBI1, NUP107, GLIPR1L2, RRN3, KXD1, HERC2, CCNT1, NOL8, RHOF, EXOSC10, CYP1B1, ST7, METTL22, ALDH18A1, SERPINH1, CTH, DGAT2, NDUFAF2, NUP54, CCDC65, CA9, CERK, GLIPR1, MARS2, MFSD8, MAST1, NFKB2, TRMT10B, REEP6, ACMSD, NDUFS3, FAM107B, BDH2, OMA1, SMCO4, C4orf33, CCDC177 |
| C-PEP.TRKB | TCF4, LRPPRC, OPRM1, EEF1A1, CARHSP1, BOLL, RCN2, CNN3, GAD2, PIK3R1, PARP12, NGFR, TRMT61A, LRRN3, TVP23B, RRN3, SUSD6, AGAP1, PLEKHM3, TMEM59L, GNA12, SLCO2B1, NPM1, HIPK2, GCDH, ACSS2, CALR, GNAQ, CNDP2, PGRMC2, DHX9, SLC45A4, REEP1, TUBGCP5, TLN1, RPS3, SLC25A44, EPS15, EHHADH, CCNT2, ELL, EFCAB5, SNIP1, GNL2, TMEM242, FAM169A, DIS3L2, NTAN1, NIPA2, LGR5, MTOR, PGS1, ACVR1, IPO9, ST7, ARL2BP, GLMP, NOCT, SLK, ZNF786, ROR1 |
| C-PEP0 | RALY, RGP1, RPS3, GADD45GIP1, C3orf18, CREB3, SMYD3, MBD2, SEPTIN4, CENPV, SCYL3, ARHGAP32, XRCC4, PEMT, RPS15, XYLB, MIER3, IER2, RRM2, POLN, DDC, C18orf54, TNFRSF11B, ZNF396, TEX14, C5AR1, TCTN3, CLIC4, LGR5, IKZF1, FGD6, HSF5, ART3, PITX1, TMEM79, CLEC5A |
| N-PEP.MRGPRD/GFRA2 | HYDIN, DGKI, STXBP6, KCNC2, NKX2-8, PLCE1, TSPAN14, DDC, NAV1, PDE7B, CENPW, GRB10, SH2D4B, NEDD4L, ANKRD44, GDPD5, PLEKHH2, PRNP, ZMAT4, FHOD3, BACH2, RALY, ABHD12, SAMD12, CARHSP1, ASAP1, RYR3, BDH2, PRKCA, TMEM132D, GLIS1, MGARP, TEN1, KIFAP3, EML6, LRRN3, CISH, ALDH18A1, DIP2C, RAD23A, TK1, MGAM, CUX2, METAP2, GNAQ, CCSER2, HECW2, ACAD11, STX10, CEP78 |
| PEP-KIT | NFIX, GPR20, MYCN, CACNA1A, NRXN3, LRRTM3, IER2, MEIS3, OPRM1, SLC44A3, ITGB1, RBPMS, EML6, DNMBP, RMDN2, PAK1, TCF4, ZMAT4, PCDH18, MRC2, PYGB, PARP12, KIFAP3, ASAP1, ELL, ATXN2, NGFR, GBF1, ACSS2, FAM117A, TNKS2, FHOD3, PRKCA, POMT1, FRMPD1, TMC8, NCAN, BEND5, PPP1R15A, CLEC5A, KIAA1328, MYCBP2, TMEM59L, ELAVL4, CYFIP1, CLN5 |

| | |
|---|---|
| | , NPR2, C5AR1, FXR1, TPM2, XYLB, ARMC9, TOM1L2, SCLT1, RNF6, ZP2, NRF1, SCUBE1, ARID4A, AQP11, FBXW11, RAB3GAP1, UCK1, DGKI, IDE, EXOSC3, POLR1E, IPO9, FBXO10, SI, COPS7B, EFCAB5, NR2C1, REEP3, LARP1B, ITCH, CCNT2, UNC13B, CALR, RAP1B, SLC9B2, LGR5, ANKS4B, SBF2, TEN1, UBA5, WDR26, MEAF6, LYST, CEP135, ZNF516, TTC39A, GLIPR1, TLN1, EXOSC10, SUGP1, CERK, NCOA6, C18orf54, POLI |
| PROPRIO | HAPLN4, CALB2, ADCY8, MET, MYL2, ADM, CRYM, SEMA4A, NTNG1, DEFB119, FSTL4, FHOD3, SH3GL2, C3orf18, LRRTM3, ACOX1, MAPKAPK3, CADM2, DNAH17, TPM2, MYOF, CNTN5, SATB2, NAT8L, PIK3R1, SLC45A4, SEPTIN4, AHCYL2, TIAM1, MBP, GALR1, PAK1, FGF1, RMDN2, SMO, HECW2, OMA1, ARHGEF39, DUT, CCSER2, ARL9, FANCG, RAB3GAP1, KCTD12, TOM1L2, BEND5, PCSK4, DYNC2LI1, CUX2, TMEM59L, PIK3C3, MTMR4, SMG5, IDE, ST7, SCYL3, ACSM5, CNN3, PPM1H, VCP, GLRA1, TMEM242, MGARP, GPR139, CCNJ, COL11A1, KIFAP3, ARL2BP, SSH2, POLR1E |
| SST | IL31RA, PLCE1, STXBP6, VWC2, RYR3, ELL, TIAM1, NAV1, FGF1, CEP78, GPR139, TCF4, PPM1H, CENPW, HYDIN, RAP1B, NOL8, DGKI, CARHSP1, RIMS2, CYP1B1, MEIS3, ASAP1, PLCXD3, KCNJ3, MAPKAPK3, FAM180B, CDC42BPA, EML6, FRMPD1, GRB10, GDPD5, GLIS1, HECW2, FHOD3 |
| TRPM8 | FOXP2, LGR5, HIPK2, ANKRD44, CALB2, ACSS2, PLCXD3, SCAPER, SNRPN, CHRM5, HEMK1, NKAIN2 |
